# The *NRC0* gene cluster of sensor and helper NLR immune receptors is functionally conserved across asterid plants

**DOI:** 10.1101/2023.10.23.563533

**Authors:** Toshiyuki Sakai, Claudia Martinez-Anaya, Mauricio P Contreras, Sophien Kamoun, Chih-Hang Wu, Hiroaki Adachi

## Abstract

NLR (nucleotide-binding domain and leucine-rich repeat-containing) proteins can form complex receptor networks to confer innate immunity. NRCs are phylogenetically related nodes that function downstream of a massively expanded network of disease resistance proteins that protect against multiple plant pathogens. Here, we used phylogenomic methods to reconstruct the macroevolution of the NRC family. One of the NRCs, we termed *NRC0*, is the only family member shared across asterid plants, leading us to investigate its evolutionary history and genetic organization. In several asterid species, *NRC0* is genetically clustered to other NLRs that are phylogenetically related to NRC-dependent disease resistance genes. This prompted us to hypothesize that the ancestral state of the NRC network is an NLR helper-sensor gene cluster that was present early during asterid evolution. We validated this hypothesis by demonstrating that NRC0 is essential for the hypersensitive cell death induced by its genetically linked sensor NLR partners in four divergent asterid species: tomato, wild sweet potato, coffee and carrot. In addition, activation of a sensor NLR leads to high-order complex formation of its genetically linked NRC0 similar to other NRCs. Our findings map out contrasting evolutionary dynamics in the macroevolution of the NRC network over the last 125 million years from a functionally conserved NLR gene cluster to a massive genetically dispersed network.

**One-sentence summary:** *NRC0* gene cluster is functionally conserved across divergent asterid species and predates the massively expanded NRC network.

## INTRODUCTION

Plants have evolved an effective innate immune system that is activated by immune receptors upon sensing diverse pathogen molecules in either the extracellular or intracellular space of host cells. One such immune receptor is the nucleotide-binding domain and leucine-rich repeat-containing protein (NLR) family, composed of intracellular immune receptors recognizing pathogen-secreted proteins called effectors (Jones et al., 2016). The evolution of NLRs is marked by a continuous arms race between plants and pathogens, leading to the rapid evolution and diversification of NLR genes even at the intraspecific level (Van de Weyer et al., 2019; Lee and Chae, 2020; Prigozhin and Krasileva, 2021). NLRs are known as the most diverse protein family in flowering plants, as many plants have hundreds to thousands of diverse NLR genes in their genomes (Shao et al., 2016; Baggs et al., 2017; Kourelis et al., 2021). This diversification and expansion of NLR genes in plants possibly occurred through genome rearrangements such as gene duplication, deletion and conversion, as well as point mutations and domain insertions (Barragan and Weigel, 2021). As a consequence of these events, NLR genes in plant genomes are often found in clusters. For example, in *Arabidopsis thaliana* accessions, 47 to 71% of NLR genes form NLR gene clusters in their genomes (Van de Weyer et al., 2019). Clustered NLR genes tend to have higher nucleotide sequence diversity than non-clustered NLR genes in Arabidopsis (Van de Weyer et al., 2019), thereby providing a reservoir of genetic variation for novel immune specificities against fast-evolving pathogen effectors. Understanding the macroevolutionary dynamics of NLRs across diverse plant species is crucial not only for unraveling the molecular mechanisms underlying plant immunity but also for providing genetic resources of disease resistance traits for global food security.

NLR and NLR-related proteins are key components of innate immunity and non-self recognition not only in plants but also in animals, fungi and bacteria (Jones et al., 2016; Uehling et al., 2017; Kibby et al., 2023). Plant NLRs have a shared domain architecture of a central nucleotide-binding domain with APAF-1, various R proteins, and CED-4 (NB-ARC) domain and a C-terminal leucine-rich repeat (LRR) domain (Kourelis et al., 2021). At the N-terminal region of plant NLRs, there is a variable domain that can be a coiled-coil (CC) or a toll/Interleukin-1 Receptor (TIR) domain. Based on the N-terminal domains, plant NLRs are broadly classified into four subfamilies: CC-NLRs (CNLs), G10-type CC-NLRs (CC_G10_-NLRs), RESISTANCE TO POWDERY MILDEW 8 (RPW8)-type CC-NLRs (CC_R_-NLRs or RNLs) and TIR-NLRs (TNLs) (Lee et al., 2021). Phylogenomic studies revealed that the most widely conserved CC-NLR gene across flowering plant (angiosperm) species is *HOPZ-ACTIVATED RESISTANCE1* (*ZAR1*), which was initially identified in Arabidopsis (Gong et al., 2022; Adachi et al., 2023). As there is no NLR gene genetically clustered to the *ZAR1* locus across angiosperm species, *ZAR1* is defined as a genetic singleton NLR gene throughout its evolution, and indeed the ZAR1 protein functions as a singleton NLR that doesn’t appear to require other NLRs to activate immunity (Adachi et al., 2023). Upon effector recognition through its LRR domain and partner receptor-like cytoplasmic kinases (RLCKs), activated ZAR1 forms a homo-pentameric complex called “resistosome”, that functions as a calcium ion (Ca^2+^) channel at the plasma membrane that is required for induction of the hypersensitive cell death immune response (Wang et al., 2019a; Wang et al., 2019b; Bi et al., 2021). To make a pore on the plasma membrane and cause Ca^2+^ influx, the ZAR1 resistosome exposes a funnel-shaped structure formed by its first α helix (α1 helix) of the N-terminal CC domain (Wang et al., 2019b; Bi et al., 2021). The α1 helix was defined as the “MADA motif” that is conserved in about 20% of CC-NLRs from dicot and monocot species (Adachi et al., 2019a). The MADA sequence is functionally interchangeable between dicot and monocot CC-NLRs, suggesting a conserved immune activation mechanism by MADA-type CC-NLRs across angiosperms. Indeed, upon effector perception, the MADA-type CC-NLR Sr35 in wheat forms the homo-oligomerized resistosome complex and causes Ca^2+^ influx (Förderer et al., 2022).

Although some plant NLRs operate as singletons, many plant NLRs have functionally specialized to recognize pathogen effectors (sensor NLRs) or to activate immune responses (helper NLRs, also known as executor NLRs) (Adachi et al., 2019b; Kourelis and Adachi, 2022). Interestingly, these functionally specialized NLRs often form gene pairs or clusters in plant genomes and function together in immune activation and regulation. For example, in Arabidopsis, *RESISTANCE TO RALSTONIA SOLANACEARUM 1* (*RRS1*) / *RESISTANCE TO PSEUDOMONAS SYRINGAE 4* (*RPS4*) (Narusaka et al., 2009; Sarris et al., 2015; Le Roux et al., 2015), *Chilling Sensitive 1* (*CHS1*) / *Suppressors of chs1-2, 3* (*SOC3*) and *TIR-NB 2* (*TN2*) (Zhang et al., 2017; Liang et al., 2019), *CONSTITUTIVE SHADE-AVOIDANCE 1* (*CSA1*) / *CHILLING SENSITIVE 3* (*CHS3*) (Xu et al., 2015; Yang et al., 2022), *SUPRESSOR OF NPR1, CONSTITUTIVE 1* (*SNC1*) / *SIDEKICK SNC1 1* (*SIKIC1*), *SIKIC2* and *SIKIC3* (Dong et al., 2018), form genetically linked pairs or clusters that function together in immune responses. In rice, *RESISTANCE GENE ANALOG 5* (*RGA5*) / *RGA4* and *PYRICULARIA ORYZAE RESISTANCE K-1* (*Pik-1*) / *Pik-2* are two sensor-helper NLR pairs that are found in head-to-head orientation in the genome (Césari et al., 2014; Maqbool et al., 2015; Shimizu et al., 2022; Sugihara et al., 2023).

In addition to NLR pairs, genetically dispersed NLRs often function together and form complex immune receptor networks. In solanaceous plants, CC-NLR proteins known as NLR-REQUIRED FOR CELL DEATH (NRC) function as helper NLRs (NRC-H) for multiple sensor NLRs (NRC-S) to mediate immune responses and to confer disease resistance against diverse pathogens (Wu et al., 2017). Although NRC-H and NRC-S genes are phylogenetically linked and form a hugely expanded NRC superclade, NRC-H and NRC-S genes are scattered throughout the genome of solanaceous plants (Wu et al., 2017). Recent biochemical and cell biology studies revealed that activated NRC-S induce homo-oligomerization of NRC-H, and activated NRC-H form punctate structures at the plasma membrane (Duggan et al., 2021; Contreras et al., 2023a; Contreras et al., 2023b; Ahn et al., 2023). This suggests that in NRC networks, NRC-H proteins activated by NRC-S presumably trigger immune responses at the plasma membrane, as is the case in the ZAR1 resistosome model. Consistent with this view, NRC-H in the NRC superclade have the MADA motif at their N-termini (Adachi et al., 2019a). In contrast, the N-termini of NRC-S genes have diversified and/or acquired additional extension domains prior to the CC domain (Adachi et al., 2019a; Seong et al., 2020). Based on these findings, a current evolutionary model of the NRC network is that NRC-H and NRC-S evolved from a multifunctional singleton NLR and have functionally specialized into helper and sensor NLRs throughout evolution (Adachi et al., 2019b; Adachi and Kamoun, 2022).

In a previous study, the NRC network was proposed to have evolved from a sensor–helper gene cluster (Wu et al., 2017). Outside of the asterid lineages, the Caryophyllales species *Beta vulgaris* (sugar beet) encodes one NRC helper and two NRC sensor genes that form a gene cluster (Wu et al., 2017). Therefore, the NRC superclade presumably emerged from a pair of genetically linked NLRs about 100 million years ago (mya) before the asterids and Caryophyllales lineages split. However, our knowledge of how NRC networks evolved across asterids and other Solanaceae-related plant species remains limited. In particular, the evolutionary dynamics of the NRC helpers across the asterids have not been studied in detail. Here, we show that *NRC0* is the only NRC-H family member that has remained conserved across asterid species. *NRC0* orthologs form gene pairs or clusters with gene(s) from the NRC-S clade, and are widely distributed in Cornales, campanulids and lamiids, but absent in Ericales. We experimentally validated the functional connections between NRC0 and genetically linked NRC-S (NRC0-S) in the *Nicotiana benthamiana* model system. Furthermore, activation of a tomato NRC0-S resulted in high-order complex formation of its genetically linked NRC0 similar to the oligomerization observed for other NRCs (Contreras et al., 2023a; Contreras et al. 2023b; Ahn et al., 2023). We propose that the *NRC0* sensor-helper gene cluster reflects the ancestral state of the NRC network; the *NRC0* cluster emerged early in asterid evolution and has massively expanded into immune receptor networks in the Solanaceae and related asterid species. This study highlights contrasting evolutionary dynamics between the functionally conserved *NRC0* gene cluster and the massively expanded and genetically dispersed NRC network of lamiid plants.

## RESULTS

### *NRC0* is the most conserved NRC helper gene in asterids

We hypothesized that the most conserved NRC gene across asterid species is most likely to reflect the ancestral state of the expanded NRC networks. To determine the distribution of helper NRC (NRC-H) genes across plant species, we first annotated NLR genes from reference genome databases of six representative plant species from asterids: carrot (*Daucus carota,* DCAR-), monkey flower (*Mimulus guttatus,* Migut-), coffee (*Coffea canephora,* Cc-), wild sweet potato (*Ipomoea trifida,* itf-), *Nicotiana benthamiana* (NbS-) and tomato (*Solanum lycopersicum*, Solyc-) by using the NLRtracker pipeline (Kourelis et al., 2021) (Supplemental File 1). To classify NRC superclade genes from the asterid NLRome dataset, we performed a phylogenetic analysis using the NB-ARC domain sequences of 1,661 annotated NLRs and 39 functionally validated NLRs (Figure 1A). In total, we identified 83 NRC-H genes from six plant species (one gene from carrot; 11 genes from monkey flower; 13 genes from coffee; 36 genes from wild sweet potato; 13 genes from *N. benthamiana*; nine genes from tomato) (Supplemental Figure S1). While most of the NRC-H form plant lineage specific subclades or clusters with previously defined Solanaceae (*N. benthamiana* and tomato) NRC-H subclade, there is one unique NRC-H subclade containing NRC-H sequences derived from four different plant species, carrot, coffee, wild sweet potato and tomato (Figure 1A; Supplemental Figure S1). We named the subclade NRC0, with each of the four species having one or two *NRC0* orthologous genes (DCAR_023561, Cc11_g06560, itf14g00240, itf14g00270, Solyc10g008220) (Supplemental Table S1). In contrast to the four species, *NRC0* was not found in monkey flower and *N. benthamiana*.

**Figure 1.**
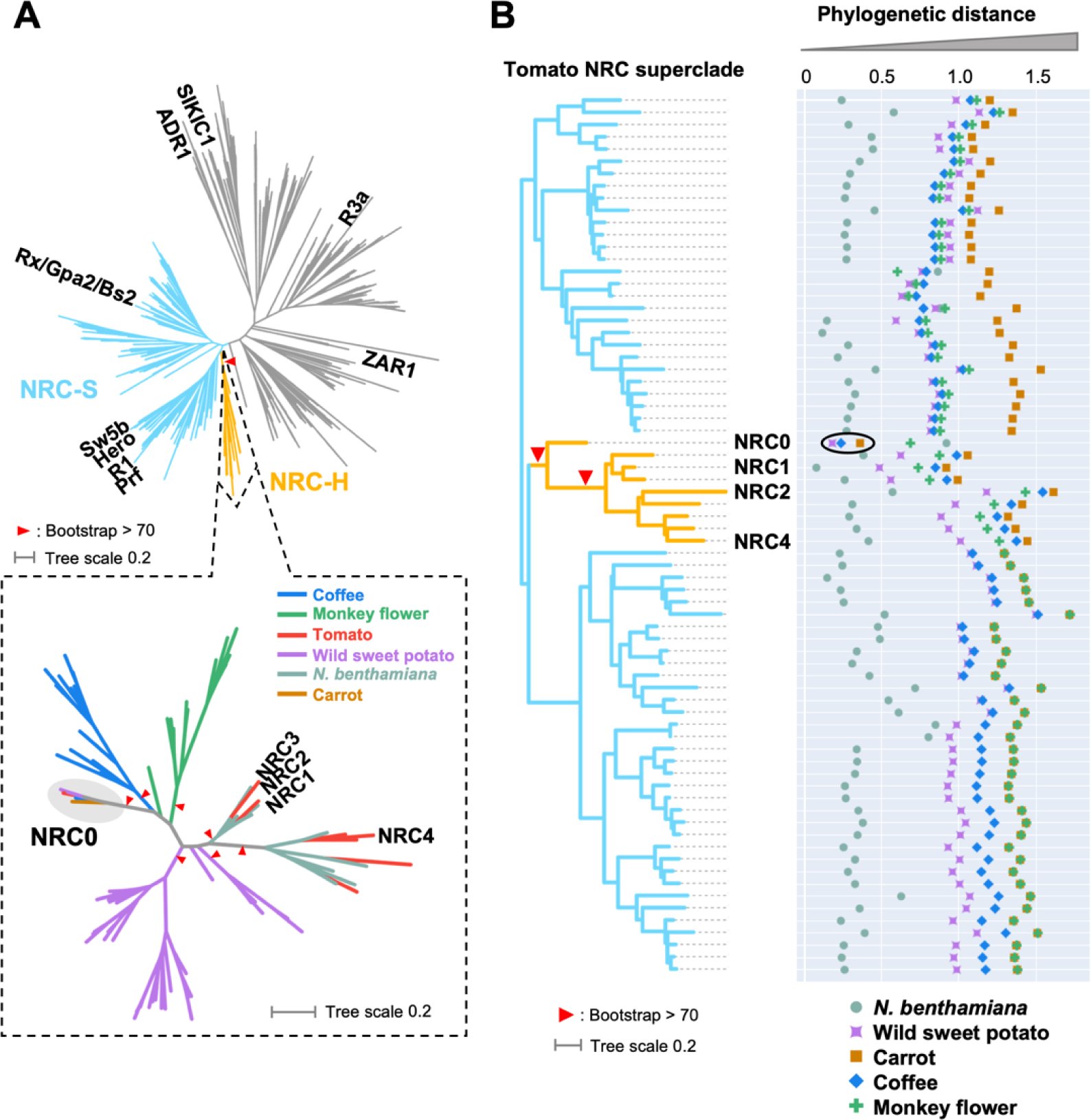
*NRC0* is the most conserved helper clade NRC in asterids. **(A)** Phylogeny of NLRs identified from asterids (carrot, monkey flower, coffee, wild sweet potato, *Nicotiana benthamiana*, and tomato). The maximum likelihood phylogenetic tree was generated in RAxML version 8.2.12 with JTT model using NB-ARC domain sequences of 1,661 NLRs identified from carrot, monkey flower, coffee, wild sweet potato, *N. benthamiana*, and tomato reference genome by using the NLRtracker and 39 functionally validated NLRs. In the top left phylogenetic tree, the NRC superclades are described with different branch color codes: NRC-helper (NRC-H) subclade shown in orange and NRC-sensor (NRC-S) subclades shown in light blue. The bottom left phylogenetic tree describes NRC-H subclade with different color codes based on plant species. Red arrow heads indicate bootstrap support > 0.7 and is shown for the relevant nodes. **(B)** Phylogenetic distance of two NRC-H and NRC-S nodes between tomato and other plant species. The phylogenetic distance was calculated from the NB-ARC phylogenetic tree shown in A. The closest distances are plotted with different colors based on plant species. Representative tomato NRC-H are highlighted.

To further evaluate *NRC0* conservation relative to other NRC-H, we used a phylogenetic tree of 805 NLR genes including NRC-H and NRC-S from the six asterid species (carrot, monkey flower, coffee, wild sweet potato, *N. benthamiana* and tomato) and nine functionally validated NRCs to calculate the phylogenetic (patristic) distance between each of the 72 tomato NRC-H/-S and their closest neighbor gene from each of the other plant species. We found that NRC0 displays the shortest patristic distance to its orthologs compared to other NRCs (Figure 1B). These phylogenetic analyses suggest that *NRC0* is possibly the most widely conserved helper NRC-H gene in asterids.

### Comparative analyses of NLR genes across asterid genomes identify a conserved *NRC0* gene cluster of candidate sensors and helpers

Plant sensor and helper NLRs often function in genetically linked pairs, while solanaceous NRCs such as NRC2, NRC3 and NRC4, form phylogenetically related but genetically dispersed NLR networks (Wu et al., 2017). To determine the degree to which helper NLRs, including *NRC0*, form NLR gene clusters in the genome, we conducted gene cluster analysis of whole NLRomes annotated from four asterid species, carrot, coffee, wild sweet potato, and tomato. In this analysis, we extracted genetically linked NLRs that have genetic distances less than 50 kb. The gene cluster information was mapped onto the NLR phylogenetic tree of the four plant species. In total, we found 1,146 genetically linked gene pairs in NLRomes of the four plant species (141 gene clusters in carrot, 324 gene clusters in coffee, 536 gene clusters in wild sweet potato, 145 gene clusters in tomato) (Supplemental Figure S2; Supplemental Table S2 and Supplemental Data Set 1).

In the NRC superclade, 438 genetically linked gene pairs exist in the genomes (three gene clusters in carrot, 45 gene clusters in coffee, 347 gene clusters in wild sweet potato, 43 gene clusters in tomato) (Figure 2A; Supplemental Table S2 and Supplemental Data Set 1). Out of 438 gene pairs, 378 are NRC-S gene pairs and 47 are NRC-H gene pairs. Notably, there are 13 genetically linked gene pairs that consist of both NRC-H and NRC-S genes (Figure 2A). Among them, only six gene pairs are conserved in the four plant species and are formed by *NRC0* and NRC-S clade genes (DCAR_023560 and DCAR_023561, Cc11_g06550 and Cc11_g06560, itf14g00240 and itf14g00250, itf14g00250 and itf14g00270, Solyc10g008220 and Solyc10g008230, Solyc10g008220 and Solyc10g008240) (Figure 2A, 2B). Although we found seven additional gene clusters (six gene clusters in sweet potato, one gene cluster in tomato), these gene clusters appeared to be plant lineage-specific (Figure 2A; Supplemental Table S2). CC_R_-NLRs, ADR1 and NRG1, are also known as conserved helper subfamily genes (Shao et al., 2016; Baggs et al., 2020; Liu et al., 2021). We noted that ADR1 and NRG1 in carrot, coffee, wild sweet potato, and tomato, form gene clusters with their paralogs, but not with genes from other NLR subfamilies (Supplemental Figure S2; Supplemental Table S2 and Supplemental Data Set 1). Taken together, these findings suggest that *NRC0* stands out as a widely conserved gene cluster of candidate helper-sensor NLRs in asterids.

**Figure 2.**
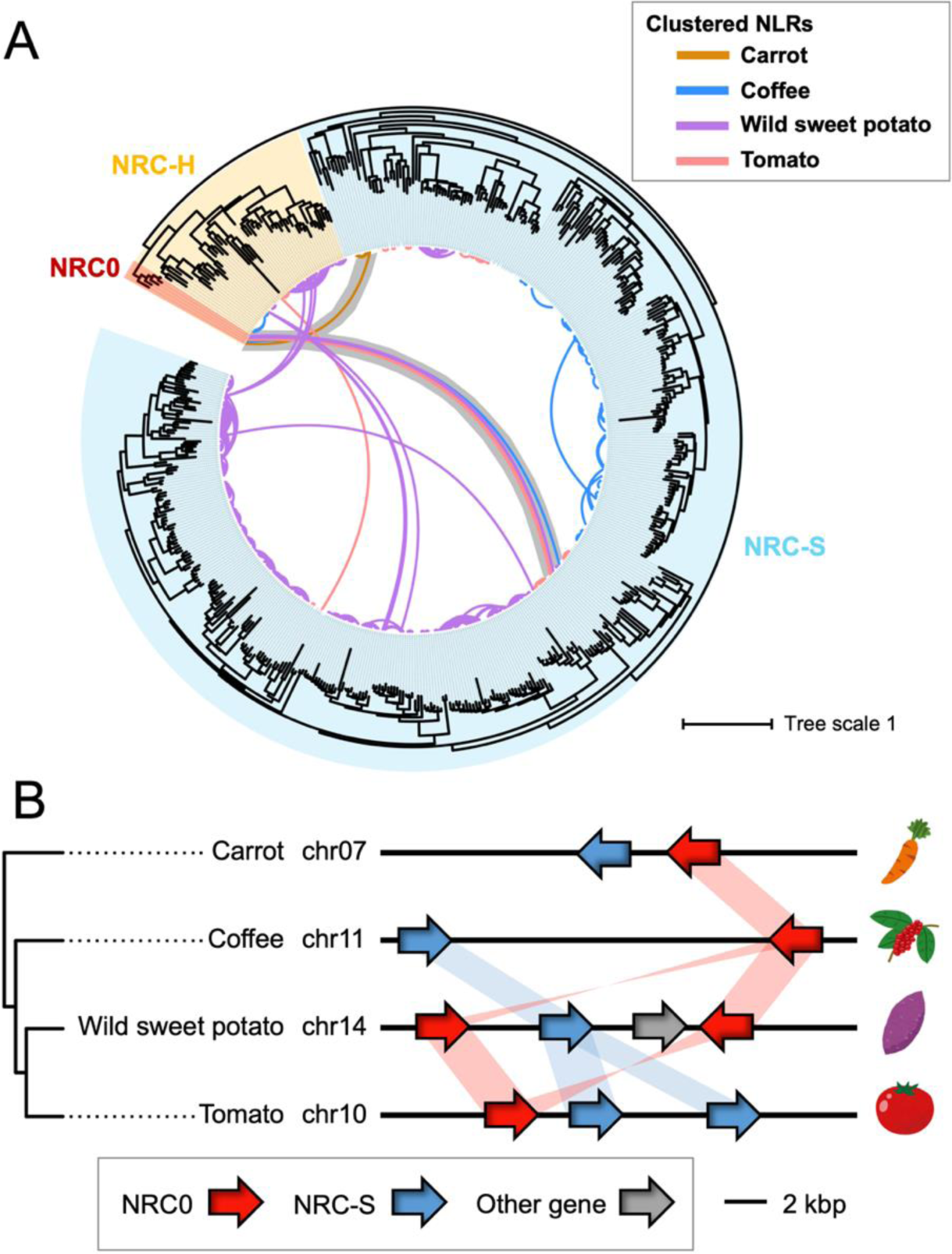
*NRC0* form a conserved gene cluster with members of the NRC sensor clade in asterids. **(A)** Phylogeny of NRC family genes from carrot, coffee, wild sweet potato and tomato with NLR gene cluster information. The maximum likelihood phylogenetic tree was generated in RAxML version 8.2.12 with JTT model using NB-ARC domain sequences of 513 NRCs identified in Figure 1. The NRC subclades are described with different background colors: NRC0 (red), other NRC-H (yellow), and NRC-S (blue). The connected lines between nodes indicate genetically linked NLRs (distance < 50 kb) with different colors based on plant species. Genetic link between *NRC0* and *NRC-S* are highlighted in gray. **(B)** Schematic representation of *NRC0* loci in carrot, coffee, wild sweet potato and tomato. Red, blue and gray arrows indicate *NRC0*, *NRC-S* genetically linked with *NRC0*, and other gene, respectively. Red and blue bands indicate phylogenetically related genes.

### The *NRC0* gene cluster predates the NRC expansion in lamiid*s*

To further examine the distribution of the *NRC0* gene cluster across plant species, we searched for *NRC0* orthologs by running a two-stage computational pipeline based on iterated BLAST searches of plant genome and protein databases and phylogenetic analysis (Figure 3A). First, we defined NRC-H genes as *NRC0* orthologs if the NRC-H genes belong to a phylogenetically well-supported clade with *NRC0* from carrot, coffee, wild sweet potato and tomato (Figure 3A, 3B). Based on this definition, we identified 38 *NRC0* orthologs from 26 asterid species, that classified in the NRC0 phylogenetic subclade (Figure 3A, 3B; Supplemental Table S1). In the second stage, we applied gene cluster analysis to identify NLR genes which are genetically linked to the obtained *NRC0* orthologs (Figure 3A). Among the 38 *NRC0* ortholog genes, 20 *NRC0* genes from 17 species are genetically linked with 23 NLR genes in NRC-S subclades (Figure 3A, 3B).

**Figure 3.**
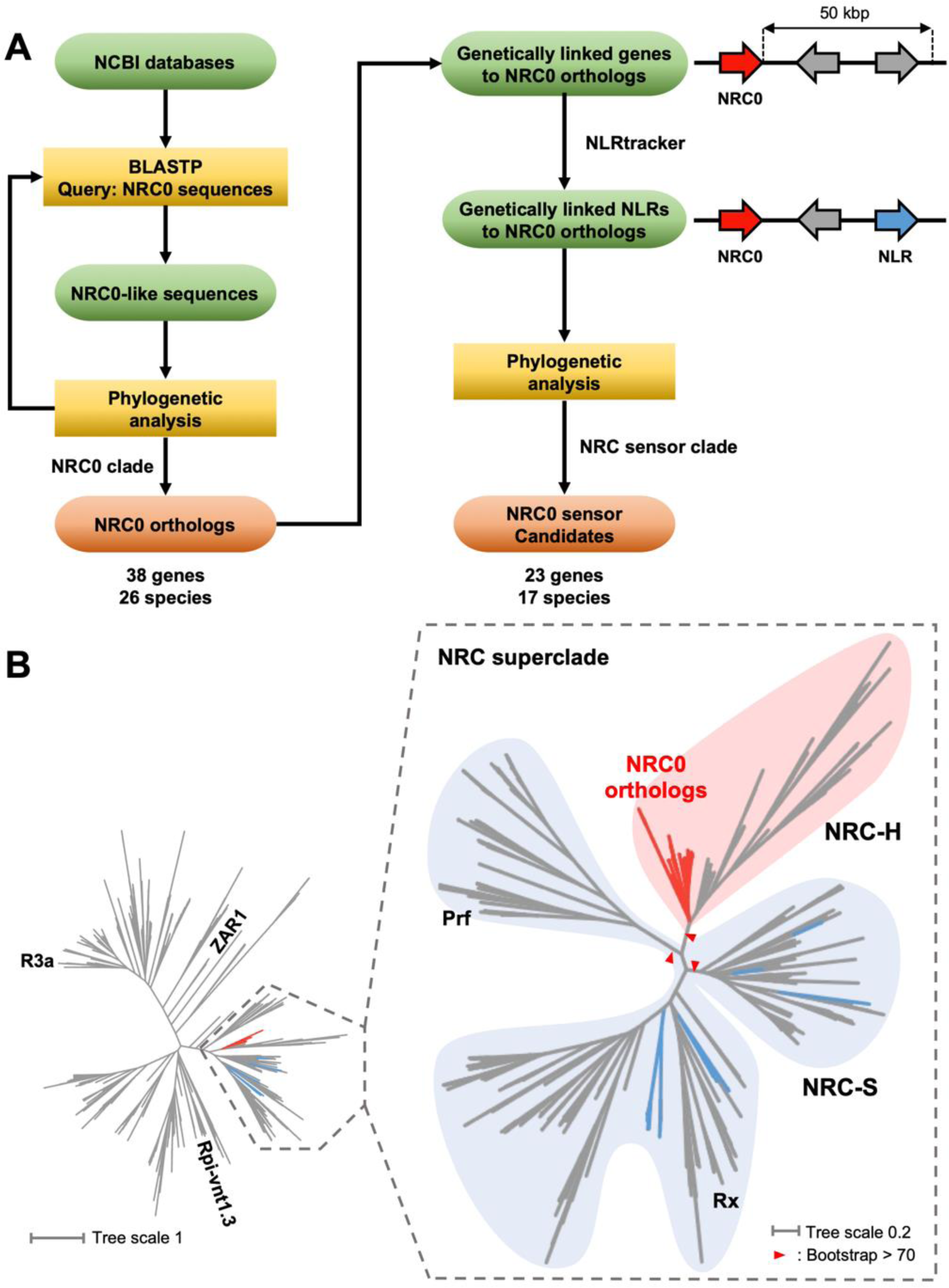
Phylogenomic analyses identify 38 *NRC0* orthologs from 26 asterid species that are linked to 23 *NRC0-S* in 17 species. **(A)** Workflow for computational analyses in searching *NRC0* orthologs and *NRC0-S* candidates. TBLASTN/BLASTP searches and subsequent phylogenetic analyses were performed to identify *NRC0* orthologs from plant genome/proteome datasets. We extracted *NRC0-S* candidates by performing gene cluster, the NLRtracker (Kourelis et al., 2021) and phylogenetic analyses. **(B)** *NRC0* orthologs exist in a subclade of the NRC-H clade. The maximum likelihood phylogenetic tree was generated in RAxML version 8.2.12 with JTT model using NB-ARC domain sequences of NRC0, NRC0-S, 15 functionally validated CC-NLRs and 1,194 CC-NLRs identified from six representative asterids, *Nyssa sinensis*, *Camellia sinensis*, *Cynara cardunculus*, *Daucus carota*, *Sesamum indicum* and *Solanum lycopersicum*. Red and blue branches indicate NRC0 and NRC0-S, respectively. Red arrow heads indicate bootstrap support > 0.7 and is shown for the relevant nodes.

Based on our criteria for the *NRC0* phylogenetic and genetic cluster, *NRC0* orthologs and their genetically linked NLRs (referred to as NRC0-dependent sensor candidates: NRC0-S) are widely distributed in asterids, Cornales, campanulids, and lamiids, but absent in Ericales (Figure 4A; Supplemental Table S1). Overall, these results suggest that the *NRC0* gene cluster emerged early in the asterid lineage.

**Figure 4.**
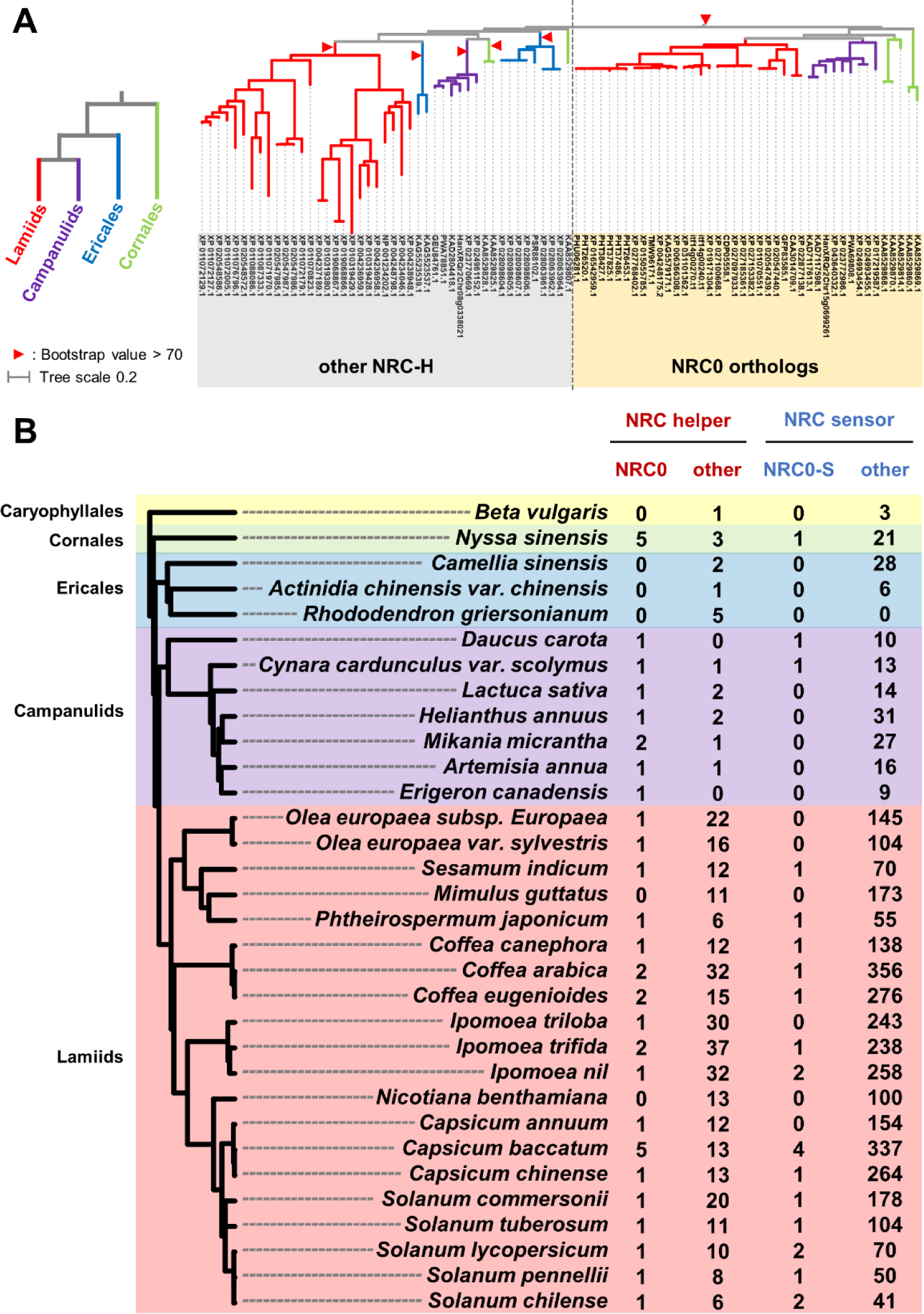
The *NRC0* gene cluster predates the massively expanded NRC network of lamiids. **(A)** Phylogeny of NRC helper subfamily defines *NRC0* orthologs and other NRCs. The maximum likelihood phylogenetic tree was generated in RAxML version 8.2.12 with JTT model using full length amino acid sequences of 87 NRC-H. The phylogenetically well-supported clade (bootstrap value > 70) containing NRC0 from Cornales, campanulids and lamiids are defined as the NRC0 subclade. The plant orders are described with different branch colors. **(B)** Distribution of the number of *NRC* genes across asterids. The left phylogenetic tree of plant species was extracted from a previous study (Smith and Brown, 2018). The right table indicates the number of *NRC0*, other *NRC-H*, *NRC0-S* and other *NRC-S* genes from 31 asterids and one Caryophyllales species. Background colors indicate plant orders.

In addition to the gene distribution analysis of *NRC0*, we investigated the number of other NRC-H and NRC-S genes across asterid species. We used NLRtracker to annotate NLR genes from 31 asterid species and one Caryophyllales species, and classified NRC-H and NRC-S genes based on phylogenetic analysis. We found NRC-H and NRC-S are drastically expanded in lamiids, compared to other plant orders in asterids (Figure 4B; Supplemental Table S3 and Supplemental File 12). In Cornales, Ericales, and campanulids, the number of NRC-H genes range from 1 to 8 and that of NRC-S genes range from 0 to 31 across species (Figure 4B; Supplemental Table S3 and Supplemental File 12). In lamiids, the number of NRC-H and NRC-S genes range from 7 to 39 and from 43 to 357, respectively (Figure 4B; Supplemental Table S3 and Supplemental File 12). Taken together, the expansion of NRC genes occurred primarily in lamiids species, but not in other asterid plants.

### *NRC0* orthologs carry the N-terminal sequence pattern required for cell death response

In the Solanaceae NRC network, the MADA motif remains at the very N terminus of NRC-H, whereas NRC-S have distinct sequences at their N-terminal region (Adachi et al., 2019a). Therefore, we hypothesized that NRC0 orthologs carry the MADA motif at their N termini for induction of cell death responses. To test this, we first ran MEME (Multiple EM for Motif Elicitation; Bailey and Elkan 1994) to search for conserved sequence patterns among the 38 NRC0 orthologs and 23 NRC0-S, respectively. All NRC0 and 10 NRC0-S carry typical CC-NB-LRR domain architecture, while 13 NRC0-S lack either CC or LRR domains (Supplemental Table S1). None of the NRC0 or NRC0-S were annotated with integrated domains that can be found in sensor NLRs for effector recognition (Figure 5; Supplemental Table S1). MEME analysis revealed 20 conserved sequence motifs that span across the NRC0 orthologs and 20 conserved sequence motifs that span across NRC0-S (Figure 5; Supplemental Table S4 and S5). Within the MEME motifs, 2^nd^ and 9^th^ MEME in the NB-ARC domain of NRC0 and 2^nd^ and 8^th^ MEME in the NB-ARC domain of NRC0-S match to p-loop and MHD motifs that coordinate binding and hydrolysis of ATP (Figure 5). Notably, in NRC0 and NRC0-S, we detected a MEME motif that is positioned at the very N-terminus where the MADA motif is generally found (Figure 5). Next, we used the HMMER software (Eddy 1998) to query the NRC0 orthologs and NRC0-S with a previously developed MADA motif–hidden Markov model (MADA-HMM; Adachi et al. 2019a). This HMMER search detected the MADA motif in 89.5% (34/38) of NRC0 orthologs, but in none of the NRC0-S (0/23) (Supplemental Table S1). Indeed, the sequence patterns of the N-termini are different between NRC0 and NRC0-S. Instead of the MADA sequence, NRC0-S have “MAHAAVVSLxQKLxx” sequence at their N-termini (Figure 5).

**Figure 5.**
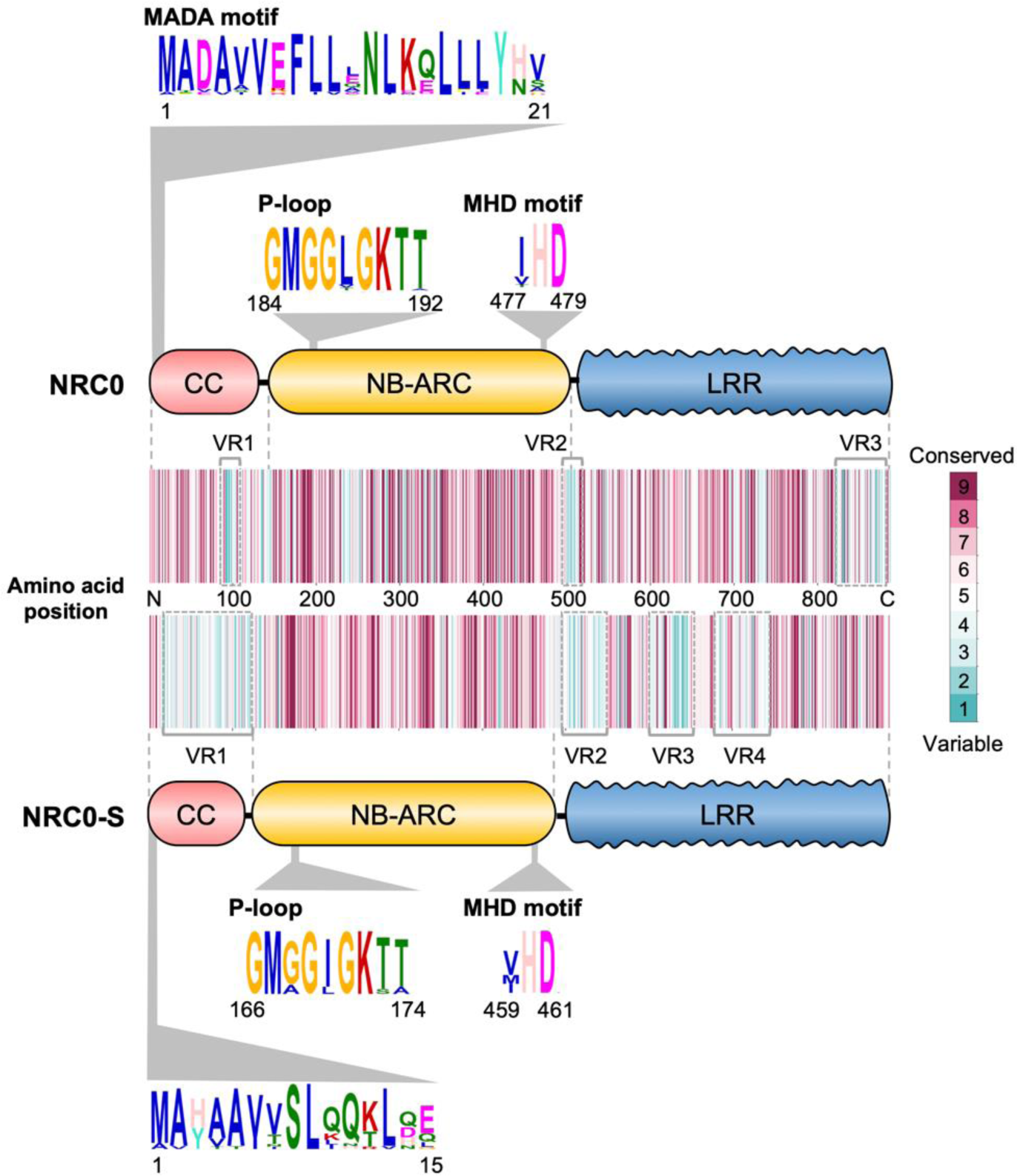
NRC0 orthologs, but not their genetically linked sensors, carry the N-terminal MADA motif required for hypersensitive cell death response. Schematic representation of conserved sequence patterns across NRC0 orthologs and NRC0 sensor candidates (NRC0-S). Consensus sequence patterns were identified by MEME using amino acid sequences of 38 NRC0 orthologs and 23 NRC0-S, respectively. Conservation and variation of each amino acid among NRC0 orthologs and NRC0-S were calculated on amino acid alignment via the ConSurf server (https://consurf.tau.ac.il). The conservation scores are mapped onto each amino acid position in tomato NRC0 (XP_004248175.2) and tomato NRC0-S (XP_004248174.1).

To further investigate sequence conservation and variation among NRC0 and NRC0-S, we used ConSurf (Ashkenazy et al., 2016) to calculate a conservation score for each amino acid and generate a diversity barcode for NRC0 and NRC0-S, respectively (Figure 5). NRC0 orthologs have highly conserved amino acid sequences across their entire domains and display few variable regions (VR) at the middle of CC domain (VR1), junction of NB-ARC and LRR domains (VR2) and C-terminal end of LRR domain (VR3) (Figure 5). In the case of NRC0-S, the amino acid sequence is varied in the CC and LRR domains (VR1 ∼ V4), although the NB-ARC domain sequence is highly conserved across NRC0-S (Figure 5).

Taken together, NRC0 orthologs carry highly conserved sequences throughout the full protein and display a canonical N-terminal MADA motif. In contrast, NRC0-S tend to be more variable especially in the CC and in parts of the LRR domains and lack a typical MADA motif.

### NRC0 are helper NLRs functionally connected with their genetically linked NLR sensors

Since NRC0, but not NRC0-S, carry the MADA motif at their N-termini, we hypothesized that NRC0 functions as a helper and NRC0-S relies on its genetic partner NRC0 to trigger immune responses. To experimentally validate this hypothesis, we first cloned *NRC0* and *NRC0-S* genes from carrot (DcNRC0: DCAR_023561, DcNRC0-S: DCAR_023560), coffee (CcNRC0: Cc11_g06560, CaNRC0-S: XM_027242939.1 in *Coffea arabica*, orthologous gene to Cc11_g06550), wild sweet potato (ItNRC0a: itf14g00240, ItNRC0b: itf14g00270, ItNRC0-S: itf14g00250) and tomato (SlNRC0: Solyc10g008220, SlNRC0-Sa: Solyc10g008230, SlNRC0-Sb: Solyc10g008240) as wild-type sequences (referred to as NRC0^WT^ or NRC0-S^WT^) (Figure 6A). We introduced an aspartic acid (D) to valine (V) mutation in the MHD motif to generate autoactive mutants of each NLR (referred to as NRC0^DV^ or NRC0-S^DV^) and tested the autoactive cell death activity in *Nicotiana benthamiana,* a species that does not have NRC0 in its genome (Figure 4B). Interestingly, four out of five tested NRC0, DcNRC0, CcNRC0, ItNRC0b and SlNRC0 caused macroscopic cell death in *N*. *benthamiana* leaves when expressed as MHD mutants, but the NRC0-S did not (Figure 6B, 6C). As a control, we expressed the *N. benthamiana* NRC-H NRC4 MHD mutant which causes autoactive cell death (Adachi et al., 2019a) (Figure 6B, 6C). Although we occasionally observed weak cell death by expressing DcNRC0-S^DV^ and SlNRC0-Sb^DV^, there was no visible cell death when expressing the majority of the DcNRC0-S^DV^ and SlNRC0-Sb^DV^ constructs, similar to the other NRC0-S (Figure 6B, 6C). This result suggests that the MADA-type CC-NLR NRC0, but not NRC0-S, has the capacity to trigger hypersensitive cell death by itself.

**Figure 6.**
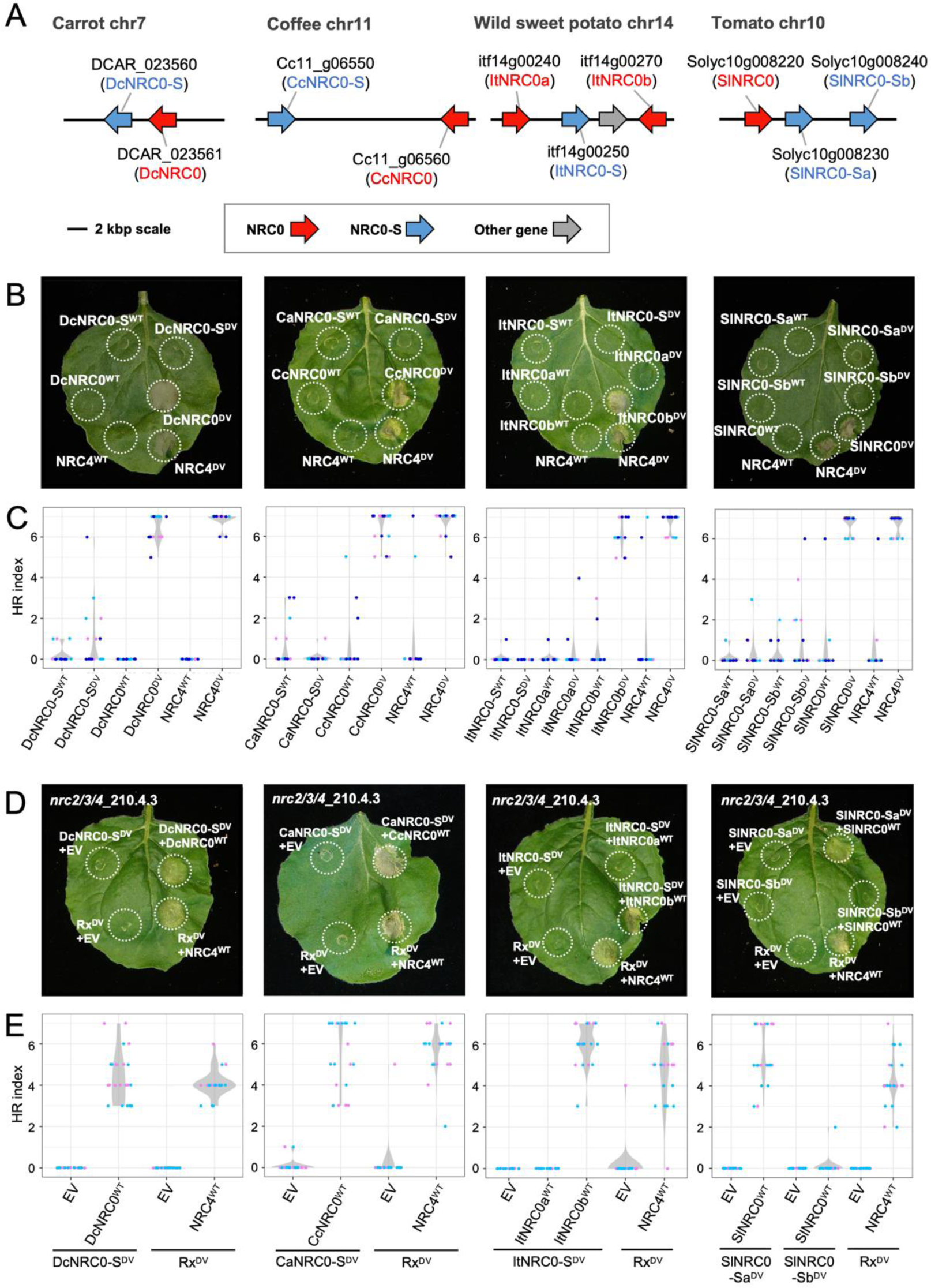
NRC0 is required for the genetically linked NRC sensor to trigger the hypersensitive cell death response in *Nicotiana benthamiana*. (A) Schematic representation of *NRC0* loci in carrot, coffee, wild sweet potato, and tomato. Red and blue arrows indicate *NRC0* and *NRC0-S*, respectively. Gray arrow indicates other gene. **(B)** Wild-type NRC0, NRC0-S, NRC4 and the MHD mutants were expressed in *N. benthamiana* leaves by agroinfiltration. Cell death phenotype was recorded five days after the agroinfiltration. **(C)** Violin plots showing cell death intensity scored as an HR index based on three independent experiments of B. **(D)** Representative images of autoactive cell death after co-expression of wild-type NRC0 (NRC0^WT^) and MHD mutants of NRC0 sensor (NRC0-S^DV^) in *N. benthamiana nrc234* mutant line. Empty vector (EV), wild-type NRC4 (NRC4^WT^) and the MHD mutant of sensor Rx (Rx^DV^) were used as controls. Photographs were taken at five days after agroinfiltration. **(E)** Violin plots showing cell death intensity scored as an HR index based on two independent experiments of D.

Our observation that *NRC0-S* are genetically clustered with helper *NRC0* prompted us to determine whether NRC0-S functionally connects with NRC0. To test this, we expressed NRC0-S MHD mutants with or without their genetically linked wild-type NRC0 in the *nrc2/3/4* knockout *N. benthamiana* line. Notably, we observed that some NRC0-S MHD mutants showed macroscopic cell death in the presence of their genetically linked NRC0 (Figure 6D, 6E). For instance, co-expression of DcNRC0-S^DV^ and DcNRC0^WT^, CaNRC0-S^DV^ and CcNRC0^WT^, ItNRC0-S^DV^ and ItNRC0b^WT^, SlNRC0-Sa^DV^ and SlNRC0^WT^ triggered a cell death response (Figure 6D, 6E). In this experiment, Rx was used as a control of NRC-dependent sensor NLR functioning with NRC-H NRC2, NRC3 and NRC4 (Wu et al., 2019). NRC4 expression complemented cell death response triggered by an autoactive MHD mutant of Rx (Rx^D460V^; Bendahmane et al., 2002) in the *nrc2/3/4* knockout line (Figure 6D, 6E). Our results indicate that NRC0-S require the genetically linked NRC0 to trigger immune response.

### Co-expression of mis-matched NRC0 and NRC0-dependent sensor pairs from different asterid species reveals evolutionary divergence

Our finding that the *NRC0* cluster is conserved in asterid species suggested that NRC0 and NRC0-S are functionally paired across asterids. However, the degree to which co-evolution between sensors and helpers has resulted in functional incompatibilities over evolutionary time is unclear. We explored whether sensor-helper pairs have functionally diverged over evolutionary time by co-expressing mis-matched pairs from different species, i.e. MHD mutants of each NRC0-S with wild-type NRC0 from four asterid species (carrot, coffee, wild sweet potato, and tomato) in the *nrc2/3/4 N. benthamiana* mutant line. We observed functional connections between NRC0-S and NRC0 across asterids with different specificities (Figure 7A, 7B; Supplemental Figure S3). For instance, co-expression of either DcNRC0-S^DV^ or CaNRC0-S^DV^ and four tested NRC0, DcNRC0^WT^, CcNRC0^WT^, ItNRC0b^WT^ or SlNRC0^WT^, triggered cell death response (Figure 7A, 7B; Supplemental Figure S3). Unlike carrot and coffee NRC0-S, ItNRC0-S^DV^ and SlNRC0-Sa^DV^ triggered cell death response only with ItNRC0b^WT^ or SlNRC0^WT^ (Figure 7A, 7B; Supplemental Figure S3). As a control, we co-expressed NRC4^WT^ with DcNRC0-S^DV^, CaNRC0-S^DV^, ItNRC0-S^DV^ and SlNRC0-Sa^DV^ and 67∼84% of tested samples did not show macroscopic cell death response (Figure 7A, 7B; Supplemental Figure S3). Taken together, NRC0-S in carrot, a species of campanulids, and coffee, the early divergent lamiids, showed functional connections across all of the tested NRC0, while NRC0-S in wild sweet potato and tomato were specifically functional only with NRC0 from Solanales.

**Figure 7.**
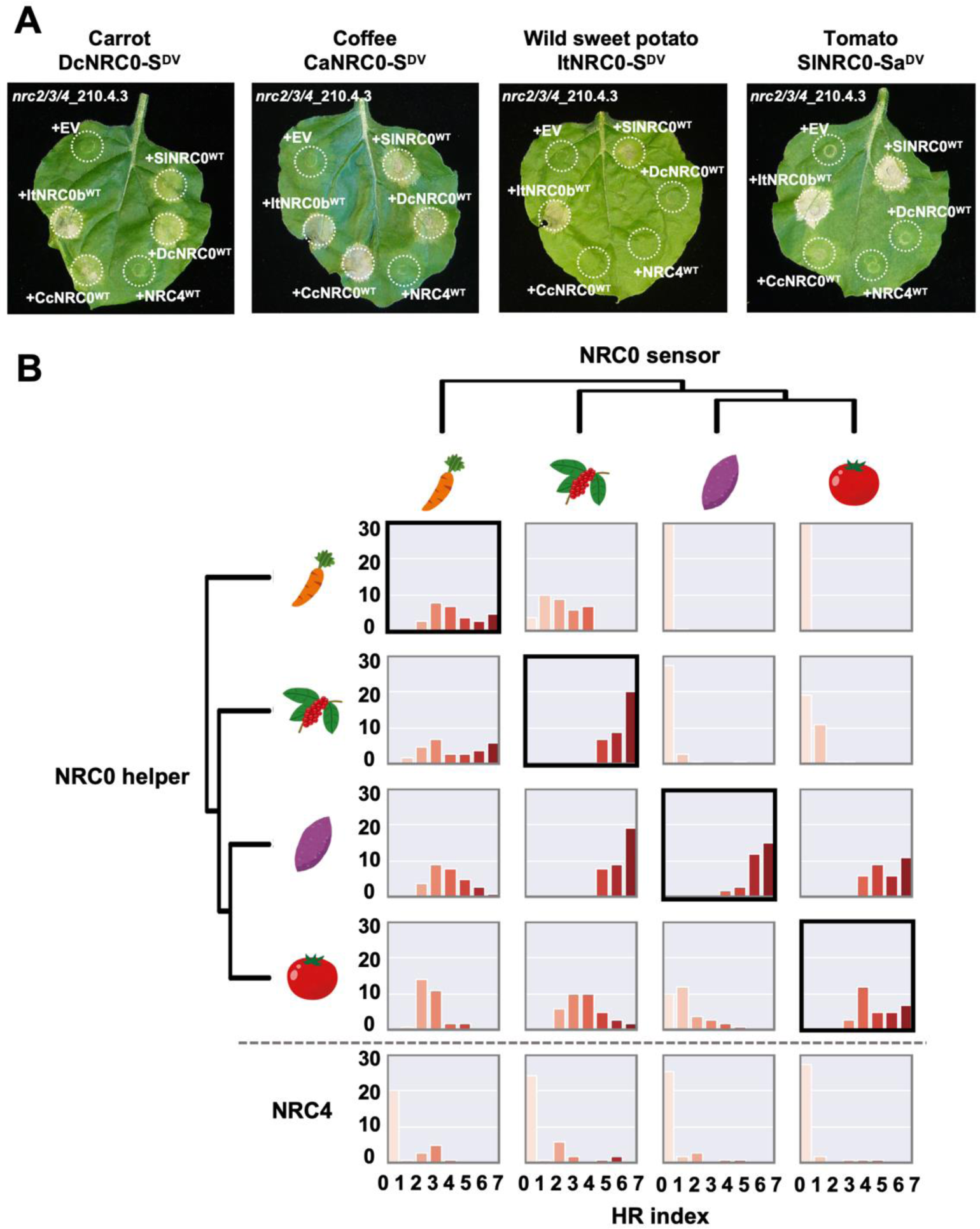
NRC0 sensors have different compatibility in inducing the hypersensitive cell death with NRC0 orthologs from across asterids. **(A)** Photographs show representative images of autoactive cell death after co-expression of MHD mutants of NRC0 sensor (NRC0-S^DV^) with wild-type NRC0 (NRC0^WT^) from four asterid species (carrot, coffee, wild sweet potato and tomato) in *N. benthamiana nrc2/3/4* mutant line. Empty vector (EV) and wild-type NRC4 (NRC4^WT^) were used as controls. Photographs were taken at five days after agroinfiltration. (B) Matrix showing the cell death response triggered by NRC0 and NRC0-S^DV^. Histograms describe cell death intensity scored in Supplemental Figure S3.

### Activated NRC0-dependent sensor leads to high-order complex formation of its genetically linked helper NRC0

Given that activation of NRC-dependent sensors induces homo-oligomerization of helper NRCs in the genetically dispersed NRC network (Contreras et al., 2023a; Ahn et al., 2023), we hypothesized that activation of NRC0-S leads to oligomerization of its genetic partner NRC0. To test this hypothesis, we first generated a MADA motif mutant of tomato NRC0 (SlNRC0^L9E/L13E/L17E^, referred to as SlNRC0^EEE^) (Figure 8A). This mutation suppresses cell death induction by MADA-type NLRs without inhibiting their resistosome formation (Adachi et al., 2019a; Hu et al., 2020; Förderer et al., 2022; Contreras et al., 2023a; Ahn et al., 2023). We expressed SlNRC0^EEE^ in *nrc2/3/4* knockout *N. benthamiana* lines with wild-type SlNRC0-Sa or its autoactive mutant SlNRC0-Sa^DV^ (Figure 8A). In inactive state with wild-type SlNRC0-Sa, SlNRC0^EEE^ was detected as a smear migrating mostly below ∼480 kDa in BN-PAGE assay (Figure 8B). Upon activation by co-expressing SlNRC0-Sa^DV^, SlNRC0^EEE^ shifted to slow-migrating higher molecular weight complex visible as a band above the 720 kDa marker (Figure 8B). This higher-order complex band of SlNRC0^EEE^ was not observed in a sample co-expressing Rx and its cognate ligand *Potato virus X* coat protein (CP), while activated Rx and CP co-expression induced oligomerization of NRC2^EEE^ in the control treatment (Contreras et al., 2023a) (Figure 8B). In this BN-PAGE assay, activated SlNRC0 showed relatively slower migration than the NRC2 oligomer (Figure 8B). This result suggests that like other NRC helpers, activated NRC0 may form the ZAR1 resistosome-type high-order complex.

**Figure 8.**
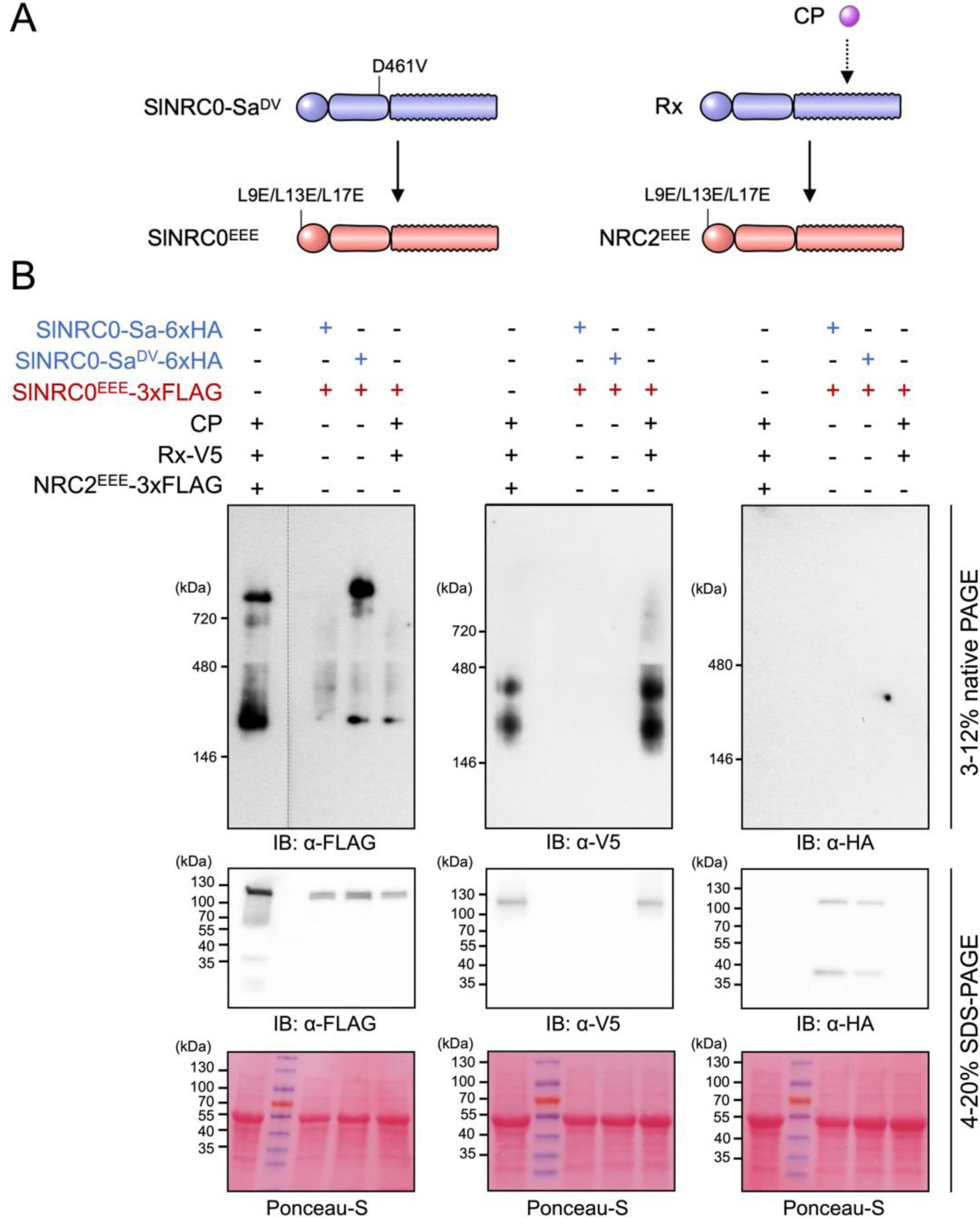
An autoactive NRC0-dependent sensor leads to formation of an NRC0 higher-order complex in *Nicotiana benthamiana*. **(A)** Schematic representation of helper NRC activation by sensor NLRs. **(B)** Detection of activated NRC0 complex in BN-PAGE. Each *Agrobacterium* strain carrying wild-type SlNRC0 sensor (SlNRC0-Sa), SlNRC0-Sa MHD mutant (SlNRC0-Sa^DV^), MADA motif mutant of SlNRC0 (SlNRC0^EEE^), *Potato virus X* coat protein (CP), wild-type Rx (Rx) or MADA motif mutant of NRC2 (NRC2^EEE^) was inoculated to leaves of an *N. benthamiana nrc2/3/4* mutant line. Total proteins were extracted from the inoculated leaves at three days after agroinfiltration. Extracts were run in native and SDS-PAGE gels, and immunoblotted with anti-FLAG, anti-V5 and anti-HA antibodies, respectively. Loading control was visualized with Ponceau-S staining. The higher-order complex of activated NRC0 was detected in three independent experiments.

Contreras et al. (2023a) and Ahn et al. (2023) showed NRC-dependent sensor NLRs are not present in the high-order complex of activated NRC, thereby proposing a model that the NRC resistosome is a homo-oligomeric complex. To investigate whether NRC0-dependent sensor NLR is associated with the activated NRC0 complex, we immunoblotted SlNRC0-Sa in the BN-PAGE assay. Although protein accumulation of wild-type SlNRC0-Sa and SlNRC0-Sa^DV^ were confirmed in an immunoblot of the SDS-PAGE assay, both signals were not detected in the BN-PAGE assay immunoblotted by anti-HA antibody (Figure 8B). In the control experiment, activated Rx appeared with two bands in the range of 146 to 480 kDa, as reported previously (Contreras et al. 2023a) (Figure 8B). In this study, we couldn’t unambiguously determine whether activated NRC0-dependent sensor integrates the resistosome complex together with its helper NRC0.

## DISCUSSION

NRC-H and NRC-S are phylogenetically related CC-NLRs that form a major superclade in asterid plants that originated from a common ancestor that predates the split between asterids and Caryophyllales (Wu et al., 2017). In solanaceous plants, the monophyletic NRC proteins function as helper NLRs for multiple sensor NLRs in a sister clade and for cell-surface localized immune receptors (Wu et al., 2017; Kourelis et al., 2022; Zhang et al., 2023). In this study, we investigated the evolutionary and functional dynamics of NRC0, an atypical member of the NRC family. *NRC0* is the only NRC family member that is conserved across asterid plants with orthologs in 26 species. *NRC0* orthologs are genetically linked to NRC-S subclade genes. We experimentally validated the functional connections within *NRC0* gene clusters for four distantly related asterid species and revealed that *NRC0* is essential for the hypersensitive cell death response triggered by its genetically linked partner *NRC0-S*. Furthermore, activated NRC0-S leads to the formation of an NRC0 high molecular weight complex similar to the model reported for other NRC-S/NRC-H pairings. We propose that the *NRC0* sensor/helper gene cluster reflects an ancestral state that predates the massive expansion of the NRC network in the lamiid lineage of asterid plants. Our findings fill a gap in the evolutionary history of an NLR network in plants and illustrates contrasting patterns of macroevolution within this complex NLR network (Figure 9).

**Figure 9.**
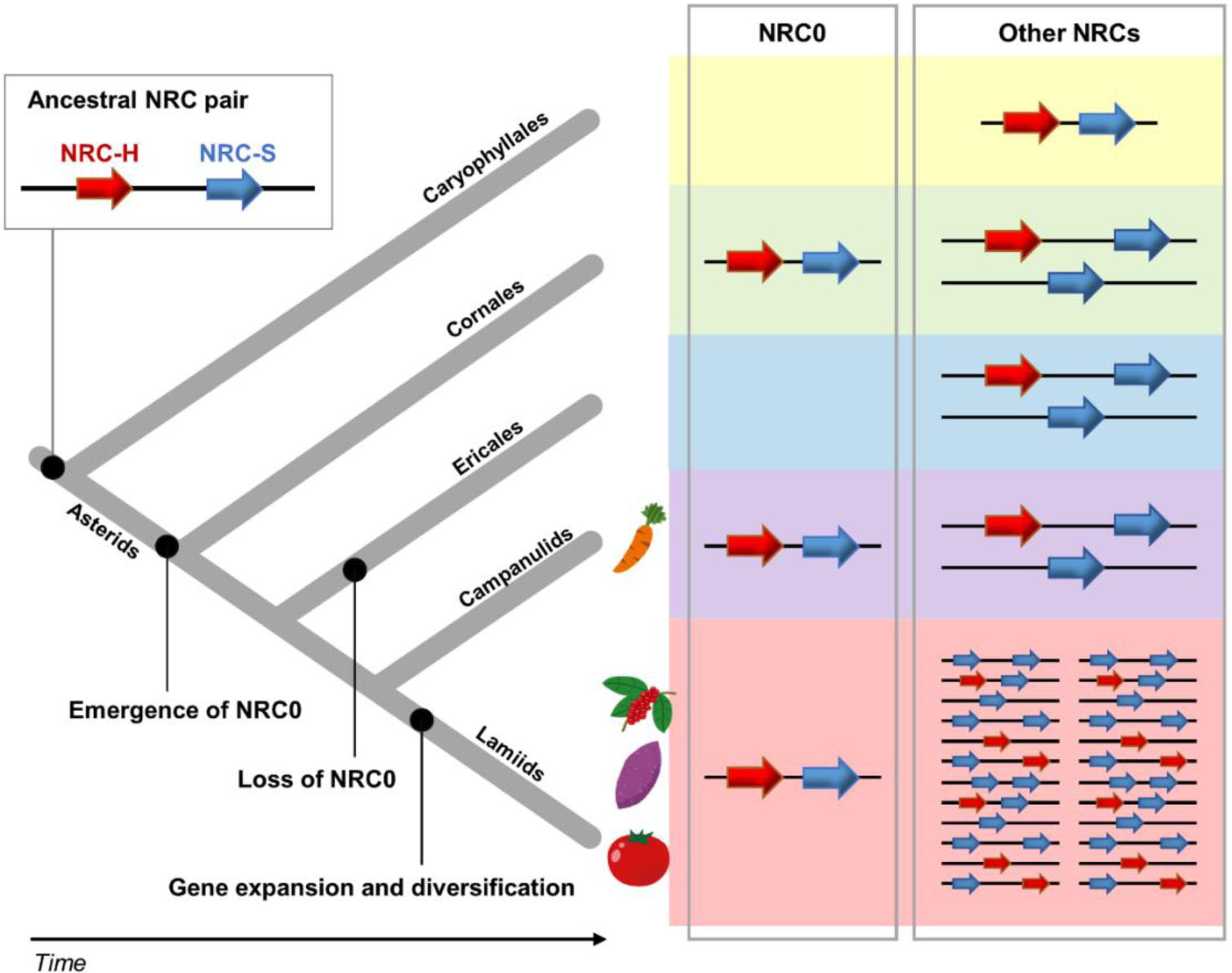
Contrasting patterns of macroevolution in the NRC network of sensor and helper NLRs. The model maps out the key evolutionary transitions in the evolution of the NRC-H and NRC-S throughout 125 million years of evolution. The NLR gene cluster of *NRC0* and *NRC0-S* has presumably originated from an ancestral NRC gene pair, which emerged before Caryophyllales and asterid lineage split. It is likely that the *NRC0* gene cluster has lost in Ericales lineage during asterid evolution. The NRC helper and sensor genes have expanded and genetically dispersed in lamiids species, while NRC components faced limited expansion in Cornales, Ericales and campanulids.

The *NRC0* gene cluster most likely emerged early in asterid evolution, which corresponds to about 125 mya based on the dating analyses of Wikström et al. (2015) (Figure 9). A previous phylogenomic study proposed that the NRC superclade expanded from a genetically linked NLR pair over 100 mya before asterids and Caryophyllales lineages split, because an NRC gene cluster exists in Caryophyllales *Beta vulgaris* (sugar beet) (Wu et al., 2017). Consistent with this, we identified one NRC-H and three NRC-S genes from *B. vulgaris*. However, the sugar beet NRC-H gene doesn’t map to the NRC0 subclade (Figure 4B). Therefore, the *NRC0* gene cluster probably originated from a common ancestral NLR pair that might be shared with the Caryophyllales NRC gene pair. We hypothesize that later during asterid evolution, the *NRC0* gene pair or a paralogous NLR gene pair have duplicated and expanded into complex NRC networks across asterid genomes (Figure 9). We noted that the expansion and diversification of NRC networks are significant in lamiids. In sharp contrast, Cornales, Ericales and campanulids have experienced limited expansions of NRC-H and NRC-S genes possibly due to low levels of NRC gene duplications and frequent deletions.

The *NRC0* gene cluster is missing in the Ericales *Camellia sinensis*, *Actinidia chinensis* var. chinensis, and *Rhododendron griersonianum*, and in some other asterid species, such as *Mimulus guttatus* and *Nicotiana benthamiana* (Figure 4B). NLRs are known to be costly genes to plants due to trade-offs between plant growth and NLR-mediated immunity and because they can cause severe autoimmune phenotypes triggered by NLR mis-regulation (Karasov et al., 2017; Adachi et al., 2019b). Thus, the *NRC0* gene cluster may have been lost as a consequence of selection against potential autoimmunity. Notably, five campanulids (*Lactuca sativa*, *Helianthus annuus*, *Mikania micrantha*, *Artemisia annua* and *Erigeron canadensis*) and four lamiids (*Olea europaea* subsp. *Europaea*, *Olea europaea* var. *sylvestris*, *Ipomoea triloba* and *Capsicum annuum*) have *NRC0* orthologs but no genetically linked *NRC0-S* encoded within 50 kb genetic distance. It is possible that *NRC0* and *NRC0-S* genes have been genetically dispersed in their genomes like in other sections of the NRC network.

Both NRC0 and NRC0-S across asterids have a typical CC-NB-LRR domain architecture. In the case of well-studied NLR pairs, Arabidopsis RRS1/RPS4, rice RGA5/RGA4 and Pik-1/Pik-2, the sensor NLRs acquired additional integrated domains that function as decoys to bait pathogen effectors (Sarris et al., 2015; Le Roux et al., 2015; Césari et al., 2014; Maqbool et al., 2015; Shimizu et al., 2022; Sugihara et al., 2023). Furthermore, in the Solanaceae NRC networks, about half of the NRC-S subclade members acquired N-terminal domain extensions that are often involved in effector recognition (Saur et al., 2015; Li et al., 2019; Adachi et al., 2019; Seong et al., 2020). Since NRC0-S do not have additional predicted domains, NRC-S likely recognize pathogen effectors through their LRR domain as is the case for the ZAR1 and Sr35 CC-NLRs (Wang et al., 2019a; Wang et al., 2019b; Förderer et al., 2022). In terms of effector perception, it is intriguing that the *NRC0* gene cluster is conserved across asterid species over 100 mya. In particular, NLRs are known to exhibit rapid evolution through a birth-and-death model (Michelmore and Meyers 1998). NRC0-S might recognize pathogen effectors in an indirect manner, either monitoring key immune signaling components of the host or functioning with other decoy components.

Our sequence motif analysis revealed that NRC0 orthologs have the MADA motif at their N-termini, but their genetically linked NRC0-S partners do not carry the canonical MADA-type sequences of CC-NLRs (Figure 5). This pattern supports the ‘use-it-or-lose-it’ model in which sensor NLRs lose the molecular signatures of the MADA motif over evolutionary time and instead rely on MADA-type helper NLRs for activation of downstream immune responses (Adachi et al., 2019a). We experimentally demonstrated that NRC0 orthologs can induce the hypersensitive cell death and are required for NRC0-S autoactive cell death (Figure 6). Although NRC0-S are not predicted to have the MADA motif and did not induce cell death without an NRC-0 helper, the “MAHAAVVSLxQKLxx” sequence is conserved at their N termini across NRC0-S proteins (Figure 5). This conservation pattern is striking because other CC domain sequences have been highly diversified among the NRC0-S (Figure 5). The N-terminal MAHA-type sequence may have a role in the molecular function of NRC0-S and was therefore maintained at their N-termini for over 100 million years.

Our BN-PAGE assays revealed that activated NRC0-S induces the formation of NRC0 high-order complexes (Figure 8). This is consistent with previous findings that activated NRC-S proteins induce formation of homo-oligomerized NRC2 resistosome (Contreras et al., 2023a; Ahn et al., 2023). In this study, we couldn’t ascertain whether activated NRC0 forms a resistosome-like high-order complex on its own or together with its sensor partner. This was presumably due to protein stability issues with SlNRC0-Sa under the BN-PAGE conditions (Figure 8). Although further biochemical studies are needed for further mechanistic insight into NRC0 activation, the current results are consistent with the activation-and-release model proposed by Contreras et al. (2023a). In the future, the NRC0 helper-sensor pairs will help map out the evolution of biochemical activation in the NRC network throughout asterid evolution.

In summary, our study helped reveal an ancestral state of the NRC network resulting in an evolutionary model in which the massively expanded NRC networks evolved from a genetically linked NLR gene pair. As illustrated in Figure 9, the NRC-type NLRs have experienced contrasting patterns of macroevolutionary dynamics over the last 125 million years from a functionally conserved NLR gene cluster to a massive genetically dispersed network. In Solanaceae, the NRC network evolved over tens of millions of years to confer resistance to pathogens and pests as diverse as viruses, bacteria, oomycetes, nematodes and insects. However, the type of pathogen effectors that are recognized by *NRC0* cluster sensor-helper pairs remains unknown. Furthermore, the structure of paired or networked NLR proteins and the determinant of functional specificities between the sensor-helper NLRs are important unanswered questions in the plant NLR research field. The NRC0 pairs will also help to map out the evolution of activation mechanisms across asterids. Future investigations that contrast the ancestral and modern states of NRC proteins will provide valuable insights into the biochemical function of NLR pairs and networks in plants and how these have evolved over 125 million years.

## Materials and Methods

### Phylogenetic analyses

For the phylogenetic analysis, we aligned NLR amino acid sequences using MAFFT v.7 (Katoh and Standley, 2013) and deleted the gaps in the alignments by our own Python script. The script is available from GitHub (https://github.com/slt666666/NRC0). NB-ARC domain sequences of the aligned NLR datasets were used for generating phylogenetic trees. The maximum likelihood phylogenetic tree was generated in RAxML version 8.2.12 with JTT model and bootstrap values based on 100 iterations. All datasets used for phylogenetic analyses are in Supplemental Files 13-18.

### Patristic distance analyses

To calculate the phylogenetic (patristic) distance, we used Python script based on DendroPy (Sukumaran and Mark, 2010). We calculated patristic distances from each NRC to the other NRCs on the phylogenetic tree (Supplemental File 13) and extracted the distances between NRCs of tomato to the closest NRC from the other plant species. The script used for the patristic distance calculation is available at GitHub (https://github.com/slt666666/NRC0).

### Gene cluster analysis

To calculate the genetic distances, we extracted gene annotation data from NCBI database as gff3 format. The genetic distances between NLR genes were calculated by our own Python script. The script is available from GitHub (https://github.com/slt666666/NRC0). We defined NLR clusters as NLR sets that genetic distances are less than 50 kb apart from each other. NLR clusters were visualized in phylogenetic tree by iTOL (Letunic and Bork., 2021). To visualize phylogenetic relationships of clustered NLRs in each specie, we developed “gene-cluster-matrix” library (https://github.com/slt666666/gene-cluster-matrix).

### NRC0 and its sensor candidate sequence retrieval

We performed BLAST (Altschul et al., 1990) using amino acid sequences of NRC0 orthologs from carrot (DCAR_023561), coffee (Cc11_g06560), wild sweet potato (itf14g00240.t1 and itf14g00270.t1), and tomato (Solyc10g008220.4.1) as queries to search NRC0-like sequences in NCBI nr or nr/nt database (https://blast.ncbi.nlm.nih.gov/Blast.cgi). In the BLAST search, we used cut-offs, percent identity ≥ 40% and query coverage ≥ 95%. NB-ARC domain sequences of the aligned sequences of the BLAST result were used for generating a phylogenetic tree and extracted NLRs located in the NRC0 clade as NRC0-like sequences. The BLAST pipeline was circulated by using the obtained sequences as new queries to search NRC0-like sequences over the angiosperm species.

We also generated a phylogenetic tree of NLR dataset of each specie, in which NRC0-like sequences were found and extracted NLRs located in the NRC0 clade as NRC0-like sequences. Finally, we generated a phylogenetic tree of NRC0-like sequences and NLR dataset of six asterid species (*Nyssa sinensis, Camellia sinensis, Cynara cardunculus, Daucus carota, Sesamum indicum, Solanum lycopersicum*) and defined NRC0 ortholog clade that include NRC0 sequences of carrot, coffee, wild sweet potato, and tomato based on phylogenetically well-supported bootstrap value. To extract *NRC0-S* gene, we extracted genes located within 50 kb from obtained *NRC0* genes and ran the NLR tracker pipeline (Kourelis et al., 2021) to annotate NLR genes among them.

### Sequence conservation analyses

Full-length amino acid sequences of the NRC0 or NRC0-S were subjected to motif searches using the MEME (Multiple EM for Motif Elicitation) (Bailey and Elkan, 1994) with parameters ‘zero or one occurrence per sequence, top twenty motifs’, to detect consensus motifs conserved in ≥90% of the input sequences. The output data are summarized in Supplemental Table S4 and S5.

To analyze amino acid sequence conservation and variation in NRC0 or NRC0-S proteins, aligned amino acid sequences of each NRC0 and NRC0-S datasets by MAFFT v.7 were used for the ConSurf pipeline (Ashkenazy et al., 2016). Tomato NRC0 (XP_004248175.2) or NRC0-Sa (XP_004248174.1) was used as a query for each analysis of NRC0 or NRC0-S, respectively. The output datasets of the ConSurf analyses are in Supplemental Data Set 2 and 3.

### Plant growth condition

Wild-type and mutant *Nicotiana benthamiana* were grown in a controlled growth chamber with temperature 22–25°C, humidity 45–65% and 16/8 hr light/dark cycle. The *NRC* knockout lines used have been previously described: *nrc2/3/4*-210.4.3 and *nrc2/3/4*-210.5.5 (Wu et al., 2020).

### Plasmid construction

Wild-type and MHD-mutant variants of *NRC0* and *NRC0-S* from tomato (*S. lycoperiscum*, Sl-), wild sweet potato (*I. trifida,* Itf-), coffee (*C. canephora* or *C. arabica,* Cc- or Ca-) and carrot (*D. carota,* DCAR-) were synthesized in pICH41155 through GENEWIZ Standard Gene Synthesis with synonymous mutations to *Bsa*I and *Bpi*I restriction enzyme sites. The synthesized genes were assembled into the binary vector pICH47732 or pICH47751 from the Golden Gate Modular Cloning (MoClo) kit (Weber et al, 2011) together with pICH51266 (35S promoter) and pICH41432 (OCS terminator) from the MoClo plant parts kit (Engler et al, 2014).

For BN-PAGE experiment, the MADA motif mutant of *SlNRC0* was amplified by Phusion High-Fidelity DNA Polymerase (Thermo Fisher) with forward mutation primer (AATGGTCTCTAATGGCTGATGCTGTTGTCGAATTTGAATTGTTAAATGAGAAACAACTAGAACTTTATCATGTGGATTTG) and reverse primer (AATGGTCTCTCGAACCAATATCTGGAGGATAGATGAC). The purified amplicon was used for Golden Gate assembly with pICSL50007 (3xFLAG) into the binary vector pICH86988 (level 1 acceptor with 35S promoter and OCS terminator). The synthetic *SlNRC0-Sa^DV^*was also assembled into the pICH86988 binary vector, together with pICSL50009 (6xHA). *Rx* and *NRC2^EEE^* constructs used were described previously (Contreras et al., 2023), and were assembled into the pJK268c vector with C-terminal tag modules pICSL50012 (V5) and pICSL50007 (3xFLAG), respectively.

Plasmid information is described in Supplemental File 19.

### Transient gene expression and cell death assays

Transient gene expression in *N. benthamiana* leaves was performed by agroinfiltration according to methods described previously (Bos et al., 2006). Briefly, four-week-old *N. benthamiana* plants were infiltrated with *Agrobacterium* Gv3101 strains carrying the binary expression plasmids. The *Agrobacterium* suspensions were prepared in infiltration buffer (10 mM MES, 10 mM MgCl_2_, and 150 μM acetosyringone, pH 5.6). To overexpress NLRs for cell death assays, the concentration of each suspension was adjusted to OD_600_ = 0.25. Macroscopic cell death phenotypes were scored according to the scale of Adachi et al. (2023).

### Protein extraction and BN-PAGE assay

Four-week-old *nrc2/3/4* plants were infiltrated with *Agrobacterium* suspensions as the concentration of each suspension is indicated in Supplemental File 19. Leaf tissue was collected at 72 hours after agroinfiltration in liquid nitrogen and was grounded in a Geno/Grinder homogenizer. Total proteins were extracted in GTMN buffer [10% glycerol, 50 mM tris-HCl (pH 7.5), 5 mM MgCl_2_, and 50 mM NaCl] supplemented with 10 mM dithiothreitol (DTT), 1x protease inhibitor cocktail (Sigma-Aldrich) and 0.2% Triton X-100 (Sigma-Aldrich), and incubated on ice for 10 min. After centrifugation at 5000 x*g* for 15 min, supernatants were used for BN-PAGE and SDS-PAGE assays.

For BN-PAGE, 25 µl of extracted protein samples were mixed with NativePAGE 5% G-250 sample additive (Invitrogen). 5 µl of each sample was loaded to NativePAGE 4 to 16% bis-tris gels (Invitrogen). Proteins were transferred to PVDF membranes with NuPAGE Transfer Buffer (Invitrogen) in a Trans-Blot Turbo transfer apparatus (Bio-Rad) by following the manufacturer’s instruction. Proteins were fixed to the membranes by incubating with 8% acetic acid for 15 min and were left to dry. For SDS-PAGE, extracted proteins were diluted in SDS loading dye and incubated at 72°C for 10 min and were loaded to 4-20% Mini-PROTEAN TGX gels (Bio-Rad). Immunoblotting was performed with antibodies: anti-hemagglutinin (HA) (3F10) HRP (Roche), anti-V5 (V2260) HRP (Roche) and anti-FLAG (M2) (Sigma) in a 1:5000 dilution.

### Accession numbers

Genome and gene information used in this study can be found from reference genome or GenBank/EMBL databases with accession numbers listed in Supplemental Data Set 5, Tables S1 and S3.

## Supplemental Data

**Supplemental Figure S1.**
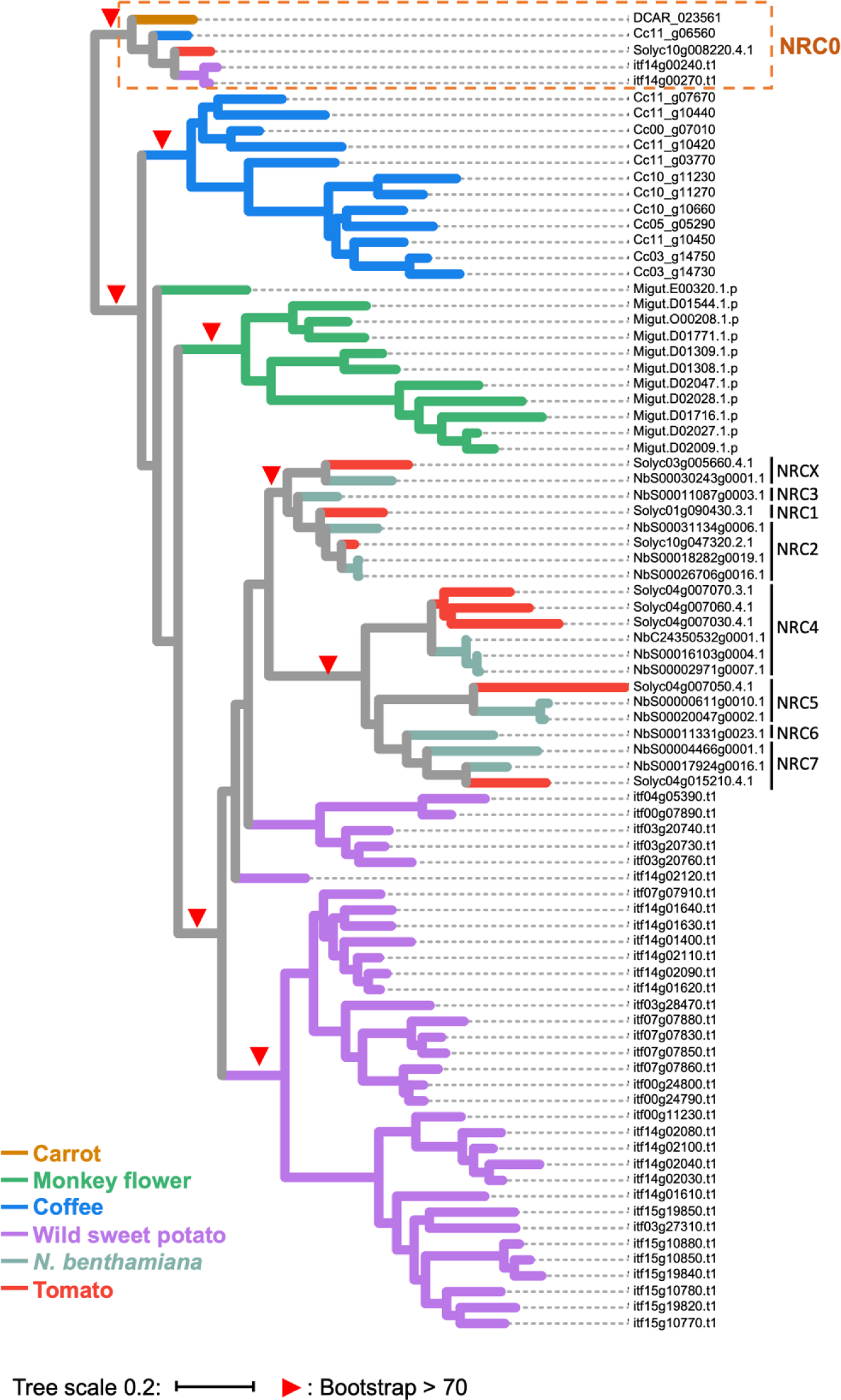
Phylogenetic tree of NRC-H clade of six asterid species. The maximum likelihood phylogenetic tree was generated in RAxML version 8.2.12 with JTT model using NB-ARC domain sequences of 83 NRC-Hs identified from carrot, monkey flower, coffee, wild sweet potato, *Nicotiana benthamiana*, and tomato. Red arrow heads indicate bootstrap support > 0.7. The scale bars indicate the evolutionary distance in amino acid substitution per site.

**Supplemental Figure S2.**
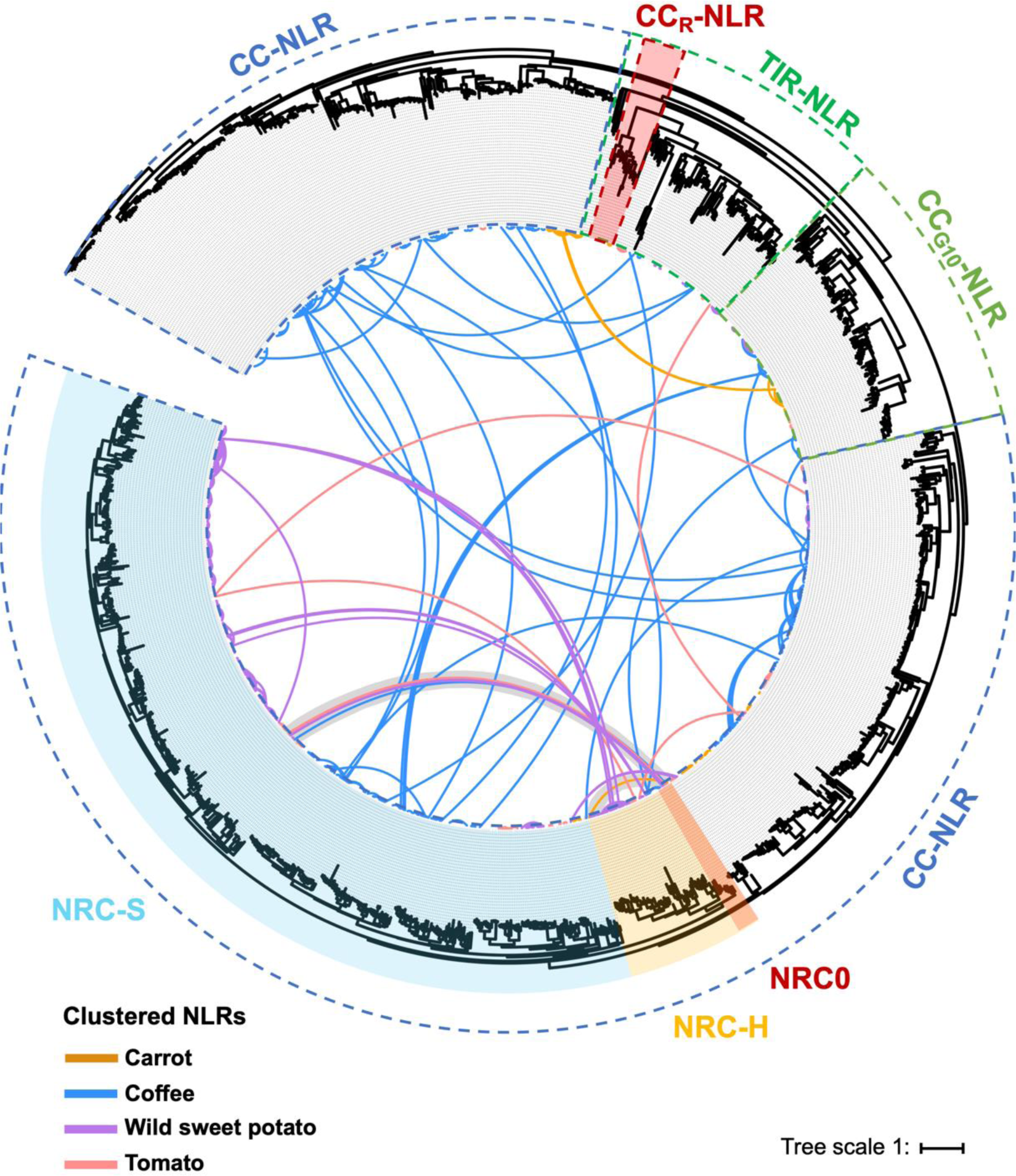
Genetically clustered NLRs in four asterid species. The maximum likelihood phylogenetic tree was generated in RAxML version 8.2.12 with JTT model using NB-ARC domain sequences of 1,265 NLRs identified from carrot, coffee, wild sweet potato and tomato. The NRC superclade containing NRC0, NRC-H, and NRC-S clades are described with different background colors. The connected lines inside the phylogenetic tree indicate genetic link between NLR genes (distance < 50 kb) with different colors based on plant species. Gray band indicate genetic link between *NRC0* and *NRC0-S*.

**Supplemental Figure S3.**
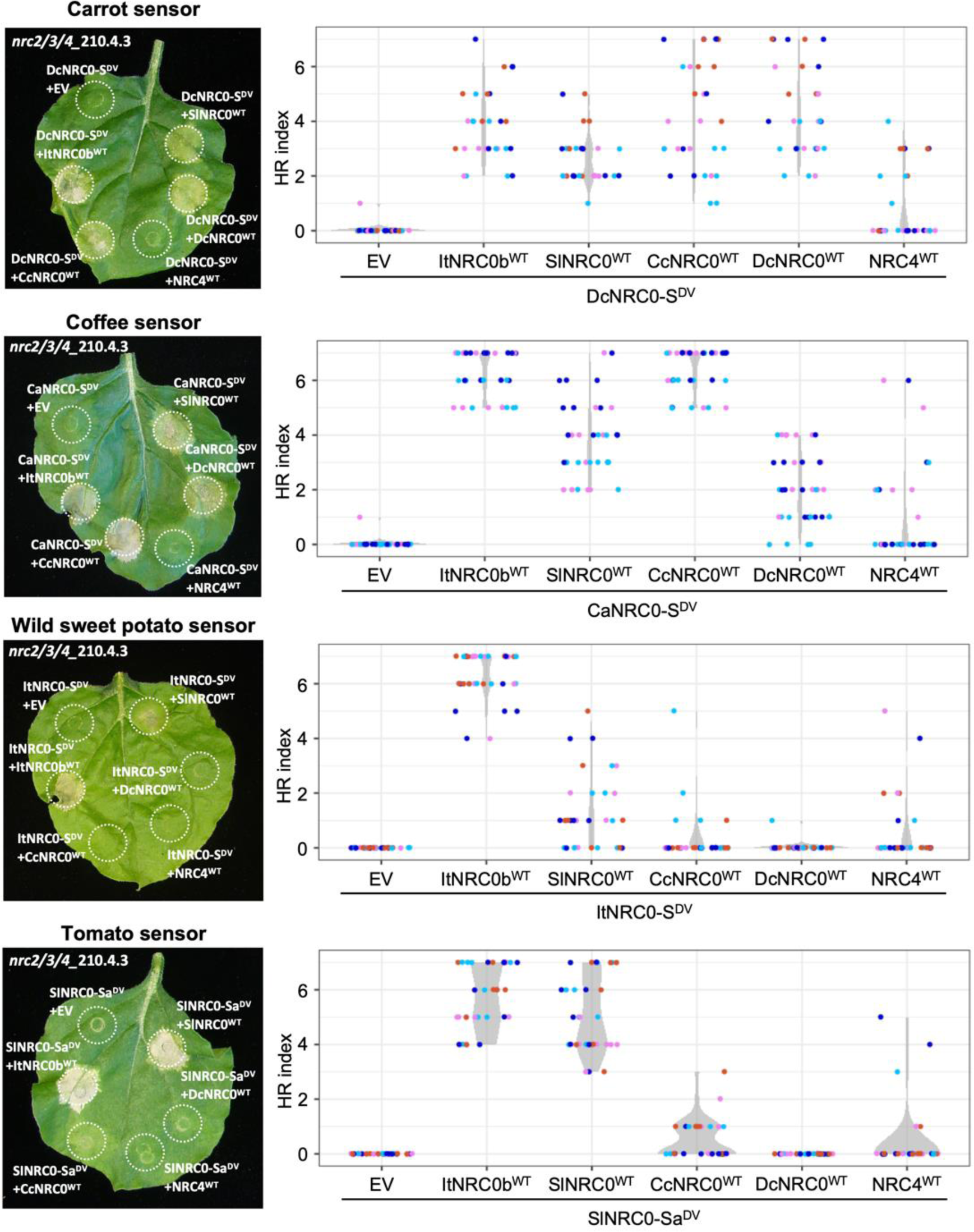
Quantification of the autoactive cell death response triggered by multiple combinations of NRC0 and NRC0-S. Left images are identical to representative images of autoactive cell death shown in Figure 7. Violin plots describe cell death intensity scored as an HR index based on three or four independent experiments.

**Supplemental Table S1.** List of *NRC0* and *NRC0-S*.

**Supplemental Table S2.** List of NLR gene cluster in four asterid species.

**Supplemental Table S3.** List of NRC in 31 asterids and 1 Caryophyllales species.

**Supplemental Table S4.** List of MEME motifs predicted from NRC0.

**Supplemental Table S5.** List of MEME motifs predicted from NRC0-S.

**Supplemental Data Set 1.** NLR gene cluster matrix files of four asterid species used for Supplemental Figure S2.

**Supplemental Data Set 2.** The ConSurf conservation score among NRC0 proteins.

**Supplemental Data Set 3.** The ConSurf conservation score among NRC0-S proteins.

**Supplemental Data Set 4.** Summary of HR index scores in cell death assay.

**Supplemental Data Set 5.** Reference genome databases used for NLR annotations.

**Supplemental File 1.** Amino acid sequences of NLRs in six asterid species and functionally validated NLRs used for Figure 1A.

**Supplemental File 2.** Amino acid alignment file of the NB-ARC domain of NLRs in six asterid species and functionally validated NLRs used for Figure 1A.

**Supplemental File 3.** Amino acid sequences of NLRs in four asterid species and functionally validated NLRs used for Figure 2A.

**Supplemental File 4.** Amino acid alignment file of the NB-ARC domain of NLRs in four asterid species and functionally validated NLRs used for Figure 2A.

**Supplemental File 5.** Amino acid sequences of CC-NLRs in six asterid species, NRC0, and NRC0-S used for Figure 3B.

**Supplemental File 6.** Amino acid alignment file of the NB-ARC domain of CC-NLRs in six asterid species, NRC0, and NRC0-S used for Figure 3B.

**Supplemental File 7.** Amino acid sequences of NRC helpers used for Figure 4A.

**Supplemental File 8.** Amino acid alignment file of full-length NRC helpers used for Figure 4A.

**Supplemental File 9.** Amino acid sequences of NRC0.

**Supplemental File 10.** Amino acid sequences of NRC0-S.

**Supplemental File 11.** Amino acid alignment file of NB-ARC domains and phylogenetic tree file of each of 31 asterid species and 1 Caryophyllales species used for Figure 4B.

**Supplemental File 12.** Amino acid sequences of NRCs in 31 asterid species and 1 Caryophyllales species.

**Supplemental File 13.** NLR phylogenetic tree file in Figure 1A.

**Supplemental File 14.** NLR phylogenetic tree file in Supplemental Figure S1.

**Supplemental File 15.** NLR phylogenetic tree file in Figure 2A.

**Supplemental File 16.** NLR phylogenetic tree file in Supplemental Figure S2.

**Supplemental File 17.** NLR phylogenetic tree file in Figure 3B.

**Supplemental File 18.** NLR phylogenetic tree file in Figure 4A.

**Supplemental File 19.** Plasmid list used in this study.

## ACKNOWLEDGEMENTS

We are thankful to our colleagues for valuable discussions and for sharing ideas. We thank Dr. Daniel Lüdke (The Sainsbury Laboratory) for valuable comments on this paper. This work was funded by the Gatsby Charitable Foundation, Biotechnology and Biological Sciences Research Council (BBSRC, UK, BB/WW002221/1, BB/V002937/1, BBS/E/J/000PR9795 and BBS/E/J/000PR9796) and the European Research Council (BLASTOFF). H.A. was funded by Japan Science and Technology Agency, Precursory Research for Embryonic Science and Technology (JPMJPR21D1). C.H.W. was funded by the 2030 Cross-Generation Young Scholars Program of the National Science and Technology Council, Taiwan (NSTC 112-2628-B-001-007). C.M.-A. is grateful to the DGAPA-PASPA UNAM Program for financing a sabbatical year at TSL.

## AUTHOR CONTRIBUTIONS

Conceptualization: S.K., C.H.W., H.A.; Data curation: T.S., C.M.-A., M.P.C.; Formal analysis: T.S., C.M.-A., M.P.C., H.A.; Investigation: T.S., C.M.-A., M.P.C., H.A.; Methodology: T.S., C.M.-A., M.P.C., H.A.; Resources: T.S., M.P.C., C.H.W., H.A.; Software: T.S.; Supervision: S.K., H.A.; Funding acquisition: S.K., H.A.; Project administration: S.K., H.A.; Writing initial draft: T.S., H.A.; Editing: T.S., C.M.-A., M.P.C., S.K., C.H.W., H.A.

