## Supplementary material for "The *NRC0* gene cluster of sensor and helper NLR immune receptors is functionally conserved across asterid plants": Supplemental Table S4.pptx

### Slide 1
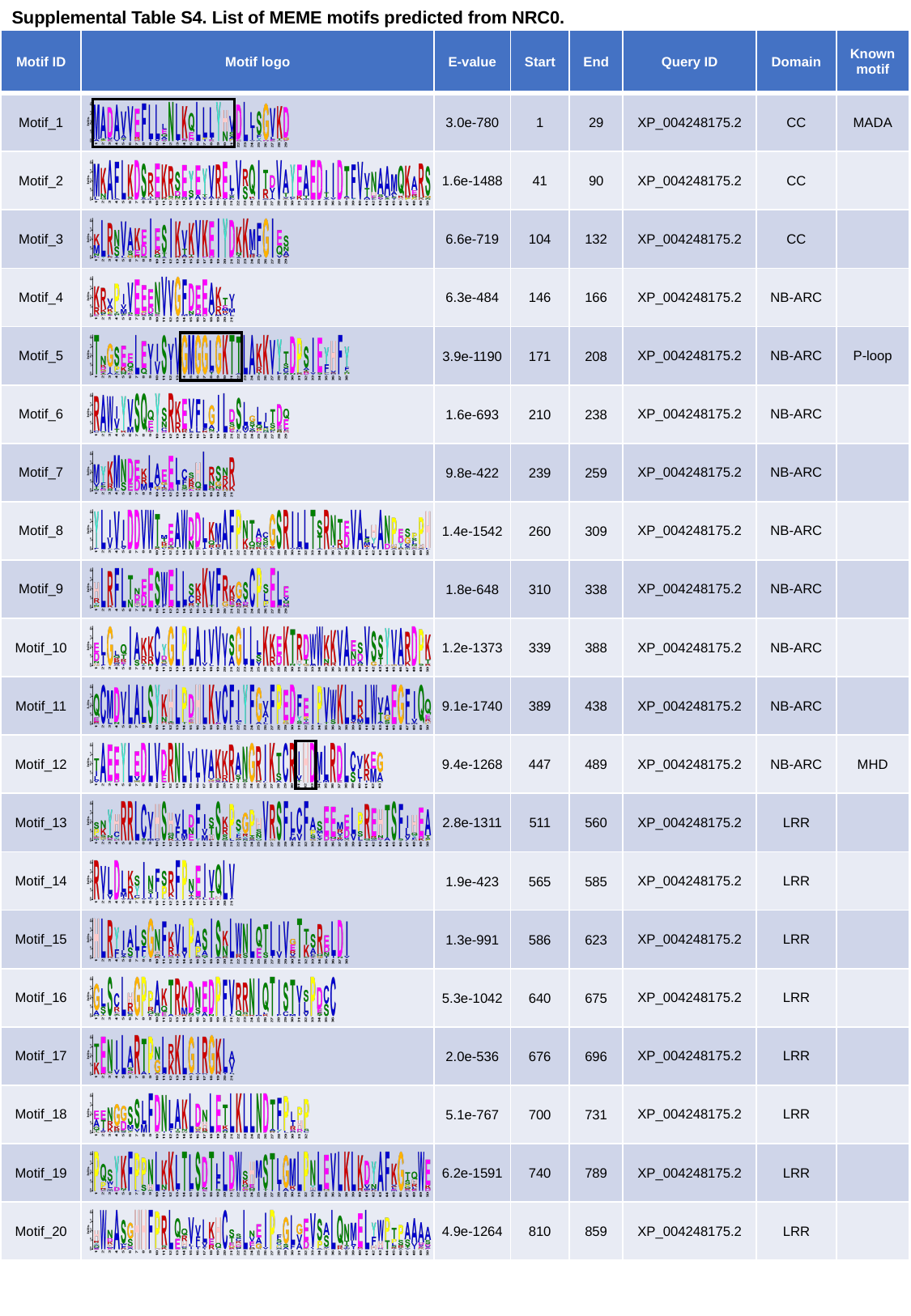

Supplemental Table S4. List of MEME motifs predicted from NRC0.
| Motif ID | Motif logo | E-value | Start | End | Query ID | Domain | Known motif |
| --- | --- | --- | --- | --- | --- | --- | --- |
| Motif\_1 | | 3.0e-780 | 1 | 29 | XP\_004248175.2 | CC | MADA |
| Motif\_2 | | 1.6e-1488 | 41 | 90 | XP\_004248175.2 | CC | |
| Motif\_3 | | 6.6e-719 | 104 | 132 | XP\_004248175.2 | CC | |
| Motif\_4 | | 6.3e-484 | 146 | 166 | XP\_004248175.2 | NB-ARC | |
| Motif\_5 | | 3.9e-1190 | 171 | 208 | XP\_004248175.2 | NB-ARC | P-loop |
| Motif\_6 | | 1.6e-693 | 210 | 238 | XP\_004248175.2 | NB-ARC | |
| Motif\_7 | | 9.8e-422 | 239 | 259 | XP\_004248175.2 | NB-ARC | |
| Motif\_8 | | 1.4e-1542 | 260 | 309 | XP\_004248175.2 | NB-ARC | |
| Motif\_9 | | 1.8e-648 | 310 | 338 | XP\_004248175.2 | NB-ARC | |
| Motif\_10 | | 1.2e-1373 | 339 | 388 | XP\_004248175.2 | NB-ARC | |
| Motif\_11 | | 9.1e-1740 | 389 | 438 | XP\_004248175.2 | NB-ARC | |
| Motif\_12 | | 9.4e-1268 | 447 | 489 | XP\_004248175.2 | NB-ARC | MHD |
| Motif\_13 | | 2.8e-1311 | 511 | 560 | XP\_004248175.2 | LRR | |
| Motif\_14 | | 1.9e-423 | 565 | 585 | XP\_004248175.2 | LRR | |
| Motif\_15 | | 1.3e-991 | 586 | 623 | XP\_004248175.2 | LRR | |
| Motif\_16 | | 5.3e-1042 | 640 | 675 | XP\_004248175.2 | LRR | |
| Motif\_17 | | 2.0e-536 | 676 | 696 | XP\_004248175.2 | LRR | |
| Motif\_18 | | 5.1e-767 | 700 | 731 | XP\_004248175.2 | LRR | |
| Motif\_19 | | 6.2e-1591 | 740 | 789 | XP\_004248175.2 | LRR | |
| Motif\_20 | | 4.9e-1264 | 810 | 859 | XP\_004248175.2 | LRR | |
