## Supplementary material for "The *NRC0* gene cluster of sensor and helper NLR immune receptors is functionally conserved across asterid plants": Supplemental Table S5.pptx

### Slide 1
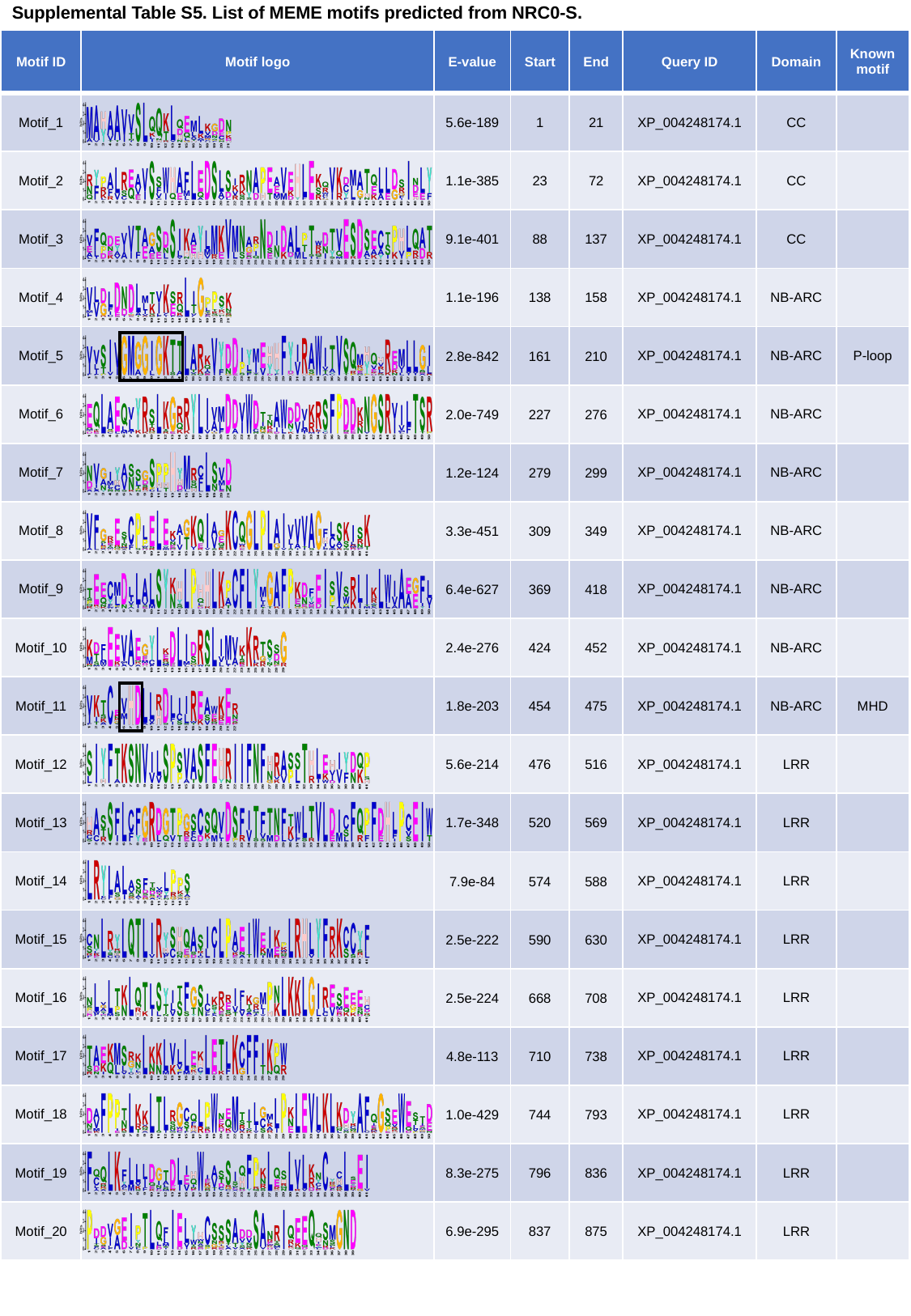

Supplemental Table S5. List of MEME motifs predicted from NRC0-S.
| Motif ID | Motif logo | E-value | Start | End | Query ID | Domain | Known motif |
| --- | --- | --- | --- | --- | --- | --- | --- |
| Motif\_1 | | 5.6e-189 | 1 | 21 | XP\_004248174.1 | CC | |
| Motif\_2 | | 1.1e-385 | 23 | 72 | XP\_004248174.1 | CC | |
| Motif\_3 | | 9.1e-401 | 88 | 137 | XP\_004248174.1 | CC | |
| Motif\_4 | | 1.1e-196 | 138 | 158 | XP\_004248174.1 | NB-ARC | |
| Motif\_5 | | 2.8e-842 | 161 | 210 | XP\_004248174.1 | NB-ARC | P-loop |
| Motif\_6 | | 2.0e-749 | 227 | 276 | XP\_004248174.1 | NB-ARC | |
| Motif\_7 | | 1.2e-124 | 279 | 299 | XP\_004248174.1 | NB-ARC | |
| Motif\_8 | | 3.3e-451 | 309 | 349 | XP\_004248174.1 | NB-ARC | |
| Motif\_9 | | 6.4e-627 | 369 | 418 | XP\_004248174.1 | NB-ARC | |
| Motif\_10 | | 2.4e-276 | 424 | 452 | XP\_004248174.1 | NB-ARC | |
| Motif\_11 | | 1.8e-203 | 454 | 475 | XP\_004248174.1 | NB-ARC | MHD |
| Motif\_12 | | 5.6e-214 | 476 | 516 | XP\_004248174.1 | LRR | |
| Motif\_13 | | 1.7e-348 | 520 | 569 | XP\_004248174.1 | LRR | |
| Motif\_14 | | 7.9e-84 | 574 | 588 | XP\_004248174.1 | LRR | |
| Motif\_15 | | 2.5e-222 | 590 | 630 | XP\_004248174.1 | LRR | |
| Motif\_16 | | 2.5e-224 | 668 | 708 | XP\_004248174.1 | LRR | |
| Motif\_17 | | 4.8e-113 | 710 | 738 | XP\_004248174.1 | LRR | |
| Motif\_18 | | 1.0e-429 | 744 | 793 | XP\_004248174.1 | LRR | |
| Motif\_19 | | 8.3e-275 | 796 | 836 | XP\_004248174.1 | LRR | |
| Motif\_20 | | 6.9e-295 | 837 | 875 | XP\_004248174.1 | LRR | |
